# Cytokinesis Arrest-induced Binucleation of Macrophages Produces Highly Efficient Efferocytes

**DOI:** 10.64898/2026.08.10.731793

**Authors:** Xun Wu, Ziyi Wang, Chenyi Xue, Fang Li, Jinnan Yue, Tyler Shern, Wenli Liu, Jian Cui, Christopher Wu, Michael Kissner, Lars Maegdefessel, Ira Tabas, Alan R. Tall, Hanrui Zhang

## Abstract

Efferocytosis, the phagocytic clearance of dying cells and debris, supports tissue homeostasis, immune tolerance, and inflammation resolution, whereas its failure contributes to autoimmunity, atherosclerosis, aging, and impaired tissue repair. Although many molecular regulators of efferocytosis have been defined, less is known about whether macrophages can be reprogrammed into a distinct cellular state with intrinsically enhanced efferocytosis capacity. Guided by a CRISPR screen, we found that *Pdcd6ip* loss induces cytokinesis arrest and binucleation, creating macrophages with superior efferocytic function. Binucleated *Pdcd6ip^−/−^* bone marrow-derived macrophages demonstrate a coordinately enhanced multi-corpse capture and processing, and resolution response, and acquired a distinct transcriptomic signature. *In vivo*, *Pdcd6ip* deletion enhanced splenic macrophage efferocytosis, reduced autoimmune responses after repeated apoptotic cell challenge, and promoted plaque stability without metabolic or hematologic changes. *PDCD6IP* perturbation similarly increased binucleation and engulfment in human macrophage-like cells. Thus, incomplete cytokinesis represents an unrecognized route to macrophage specialization with enhanced efferocytosis capacity.

## INTRODUCTION

Efferocytosis is the specialized phagocytic clearance of apoptotic cells and dying- or dead cell-derived debris by professional and non-professional phagocytes^1^. In tissues with continuous cell turnover, efficient efferocytosis prevents secondary necrosis, limits the release of intracellular danger signals, and promotes immune quiescence and tissue repair^1^. When efferocytosis fails, uncleared dying cells accumulate and amplify inflammation, autoantigen exposure, and tissue injury^1^. Dysregulated efferocytosis has therefore been implicated across diverse disease settings, including autoimmunity, atherosclerosis, cancer, age-related tissue degeneration, and impaired tissue repair after injury^2^. Accordingly, restoring defective efferocytosis provides promising therapeutic opportunities^2^.

Over the past two decades, major progress has defined molecular steps of efferocytosis, including the release of find-me signals, recognition of eat-me signals such as phosphatidylserine, apoptotic cell binding, engulfment, phagosome maturation, lysosomal degradation, and downstream resolution programs^3^. This stepwise framework involves multiple receptors and bridging molecules, including MERTK, AXL, TIM4, CD36, MFGE8, GAS6, and complement components, as well as intracellular pathways that coordinate cytoskeletal remodeling, phagolysosomal maturation, metabolic adaptation, and anti-inflammatory mediator production^3^. Yet efferocytic capacity is not uniform across macrophages^4^. *In vitro*, M2c-polarized human macrophages clear early apoptotic cells more efficiently than other macrophage subsets, in part through MERTK induction^5^. Tissue-resident TIM4-expressing macrophages are specialized for apoptotic cell capture and cooperate with TAM-family signaling pathways to promote efficient clearance^6^. In atherosclerosis, single-cell and spatial studies have identified plaque macrophage subsets enriched for TREM2, C1Q, HMOX1, CD9, lysosomal genes, and lipid-handling programs, and functional studies support roles for TREM2 and C1q in macrophage survival, lipid handling, efferocytosis, and limitation of plaque necrosis^7–9^. Together, these studies have established both molecular targets for enhancing individual steps of efferocytosis and macrophage subsets with naturally differing efferocytic capacities. An especially compelling next step is to determine whether macrophages can be deliberately reprogrammed into a distinct cellular state in which multiple stages of apoptotic cell clearance are coordinately enhanced. Such an approach could enable sustained, high-capacity clearance beyond that achieved by manipulating a single pathway and may offer broader therapeutic potential.

Multinucleated giant cells (MGCs) provide a potential clue that major changes in macrophage cellular architecture can support such functional specialization. Macrophages undergo architectural transformation when they fuse to form MGCs in granulomas, foreign-body reactions, and other chronic inflammatory contexts^10–12^. In these settings, MGCs acquire specialized capacities for large-target destruction and complement-mediated phagocytosis^13–18^. Thus, macrophage multinucleation can be associated with specialized clearance functions, but it has been studied primarily as a reactive adaptation driven by cell-cell fusion. Interestingly, oxidative damage-induced cytokinesis arrest can also promote MGC formation in granulomas^19^, revealing a non-fusion route to macrophage multinucleation. Because multinucleation is often associated with response to chronic inflammation or indigestible materials, its potential as an engineerable, beneficial cellular state remains unexplored. This leaves a critical conceptual gap: whether multinucleation can be intentionally harnessed via cytokinesis arrest to generate a unique macrophage state specialized for high-capacity clearance, and whether such a state can confer disease protection *in vivo*.

Our recent unbiased screening provided an unexpected entry point to address this gap. We performed a genome-wide CRISPR screen designed to identify regulators of efferocytosis^20^, capturing both genes required for efferocytosis^20,21^ and genes whose loss enables macrophages to continuously engulf multiple apoptotic cells. Among the top negative regulators identified was *Pdcd6ip*, which encodes PDCD6IP (Programmed Cell Death 6 Interacting Protein), also known as ALIX^22^. PDCD6IP is an adaptor protein that links the endosomal sorting complex required for transport (ESCRT) machinery to membrane remodeling events^23–25^. In addition to its established roles in multivesicular body formation^26^ and viral budding^27^, PDCD6IP has been implicated in the late stages of cytokinesis, where ALIX and ESCRT components function at the midbody and abscission site to complete daughter-cell separation^22,28–30^. This convergence between a CRISPR screen-identified efferocytosis regulator and cytokinetic abscission led us to hypothesize that *Pdcd6ip* loss could enhance efferocytosis by inducing cytokinesis failure, thereby generating a distinct binucleated macrophage state specialized for high-capacity clearance.

Here, we demonstrate that *Pdcd6ip* deletion induces binucleated macrophage formation coupled with enhancement of apoptotic cell binding, engulfment, degradation, and downstream resolution responses. Remarkably, these binucleated macrophages exhibit an exceptional capacity for multi-cargo engulfment. This phenotype is not explained solely by increased cell size, as single-cell RNA sequencing (scRNA-seq) identifies a distinct transcriptomic signature in a subpopulation of *Pdcd6ip^−/−^* macrophages enriched for binucleation. *In vivo*, *Pdcd6ip* deletion enhances apoptotic cell clearance, protects mice from autoimmune responses due to chronic apoptotic cell burden, and stabilizes atherosclerotic plaques after hematopoietic reconstitution. Together, these findings identify a link between cytokinetic abscission and macrophage efferocytic specialization, establishing PDCD6IP loss-induced binucleation as a distinct macrophage state that couples altered cellular architecture to broadly enhanced efferocytosis capacity and disease protection.

## RESULTS

### *Pdcd6ip* deletion increases macrophage efferocytosis

To determine whether PDCD6IP regulates macrophage efferocytosis, we obtained bone marrow-derived macrophages (BMDMs) from *Pdcd6ip^+/+^*and *Pdcd6ip^−/−^* mice (**Fig. 1a**, schematic of the experimental design) and confirmed loss of PDCD6IP protein by Western blotting (**Fig. 1b**). We then performed an *in vitro* efferocytosis assay using apoptotic cells (ACs) generated by UV irradiation of murine thymocytes, which were used throughout the study unless otherwise indicated. ACs were labeled with Hoechst and co-incubated with BMDMs for 45 min, after which efferocytosis was quantified by flow cytometry as the percentage of Hoechst-positive BMDMs.

**Fig. 1.**
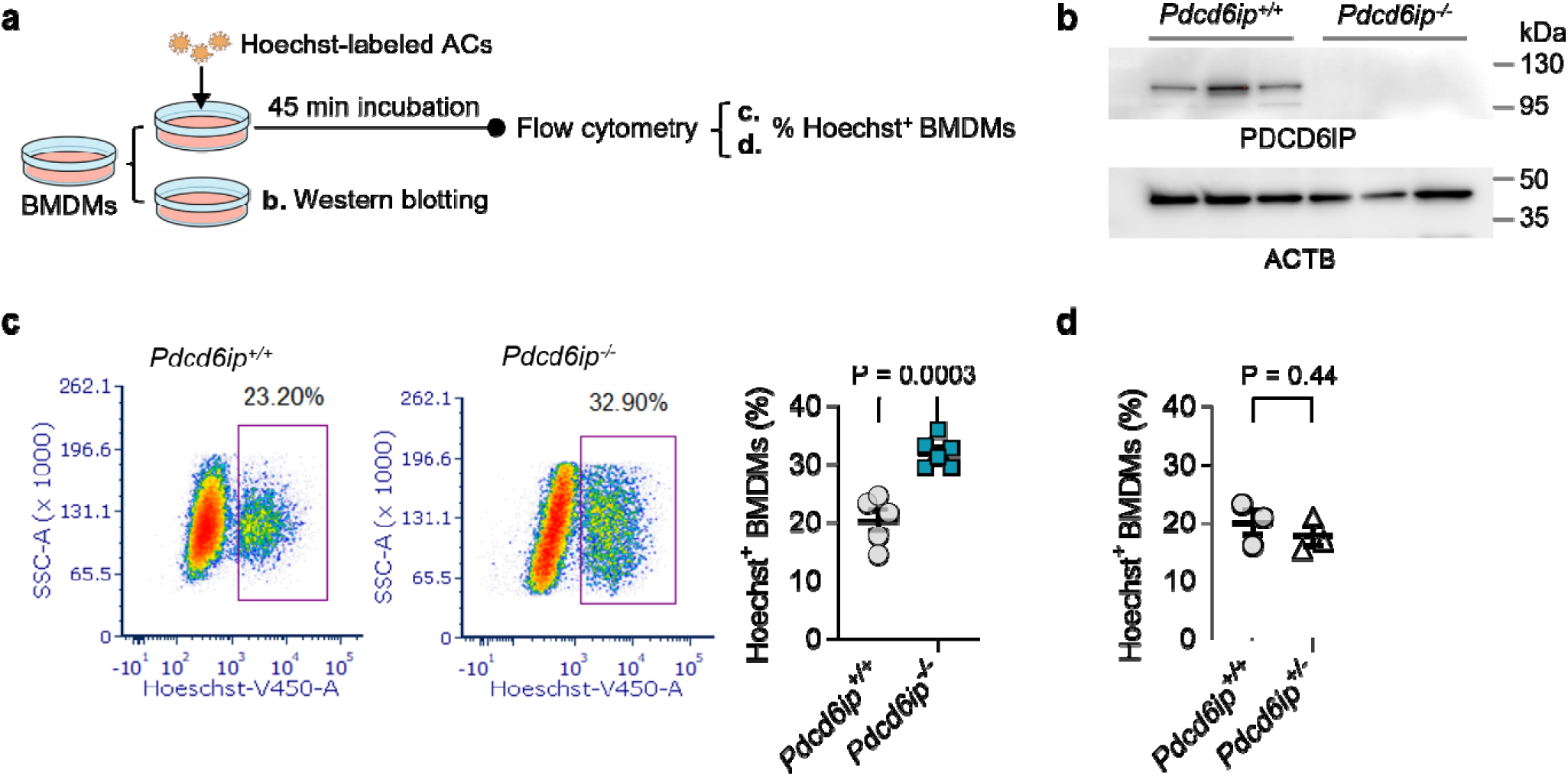
*Pdcd6ip* deletion increases macrophage efferocytosis. **a**, Schematic of the experimental design and *in vitro* efferocytosis assay. Bone marrow cells were isolated from wild-type (*Pdcd6ip*^+/+^) and knockout (*Pdcd6ip*^−/−^) mice and differentiated into bone-marrow-derived macrophages (BMDMs). Apoptotic cells (ACs) were generated by UV irradiation of mouse thymocytes, labeled with Hoechst, and co-incubated with BMDMs for 45 min. Efferocytosis was quantified by flow cytometry as the percentage of Hoechst^+^ BMDMs **b,** Western blot analysis confirming loss of PDCD6IP protein in *Pdcd6ip*^−/−^ BMDMs. n = 3 *Pdcd6ip*^+/+^ and 3 *Pdcd6ip*^−/−^ mice. **c,** Representative flow cytometry plots and quantification of Hoechst^+^ BMDMs. *Pdcd6ip* deletion enhanced efferocytosis *in vitro*. Representative flow cytometry plots showed an altered side scatter profile in *Pdcd6ip*^−/−^ BMDMs. *n* = 5 *Pdcd6ip*^+/+^ and 6 *Pdcd6ip*^−/−^ mice. **d,** Heterozygous *Pdcd6ip* deletion (*Pdcd6ip^+/^*^−^) did not enhance efferocytosis *in vitro*. *n* = 3 *Pdcd6ip^+/+^* and 3 *Pdcd6ip^+/^*^−^ mice. Data are shown as mean ± standard error of the mean (SEM). Statistical significance was determined using a two-tailed Student’s *t*-test.

*Pdcd6ip^−/−^* BMDMs showed significantly increased efferocytosis compared with *Pdcd6ip^+/+^* BMDMs (**Fig. 1c**). This phenotype was not observed in heterozygous macrophages, as *Pdcd6ip^+/−^* BMDMs exhibited efferocytosis comparable to *Pdcd6ip^+/+^* controls (**Fig. 1d**), indicating that complete loss of PDCD6IP is required to increase AC uptake under these conditions. In parallel, flow cytometry analysis revealed that *Pdcd6ip^−/−^* BMDMs had an increased side scatter, suggesting greater intracellular complexity (**Fig. 1c**). The scatter profile and the established role of PDCD6IP in cytokinetic abscission in other cell types^22,28–30^ prompted us to assess whether *Pdcd6ip* deletion alters macrophage cell architecture and nuclear content.

### *Pdcd6ip* deletion induces binucleated macrophages with enhanced apoptotic cell binding, engulfment, lysosomal acidification, and resolution responses

Microscopy-based analysis revealed a marked increase in binucleated macrophages among *Pdcd6ip^−/−^* BMDMs, with approximately 60% of *Pdcd6ip^−/−^* BMDMs containing two nuclei compared with a much smaller fraction in *Pdcd6ip^+/+^* BMDMs (**Fig. 2a**). Imaging cytometry independently confirmed this increase in binucleation (**Supplementary Fig. 1**). These observations are consistent with a role for PDCD6IP in cytokinesis completion and suggest that *Pdcd6ip* deletion causes macrophages to undergo incomplete abscission, producing a binucleated/polyploid macrophage population.

**Fig. 2.**
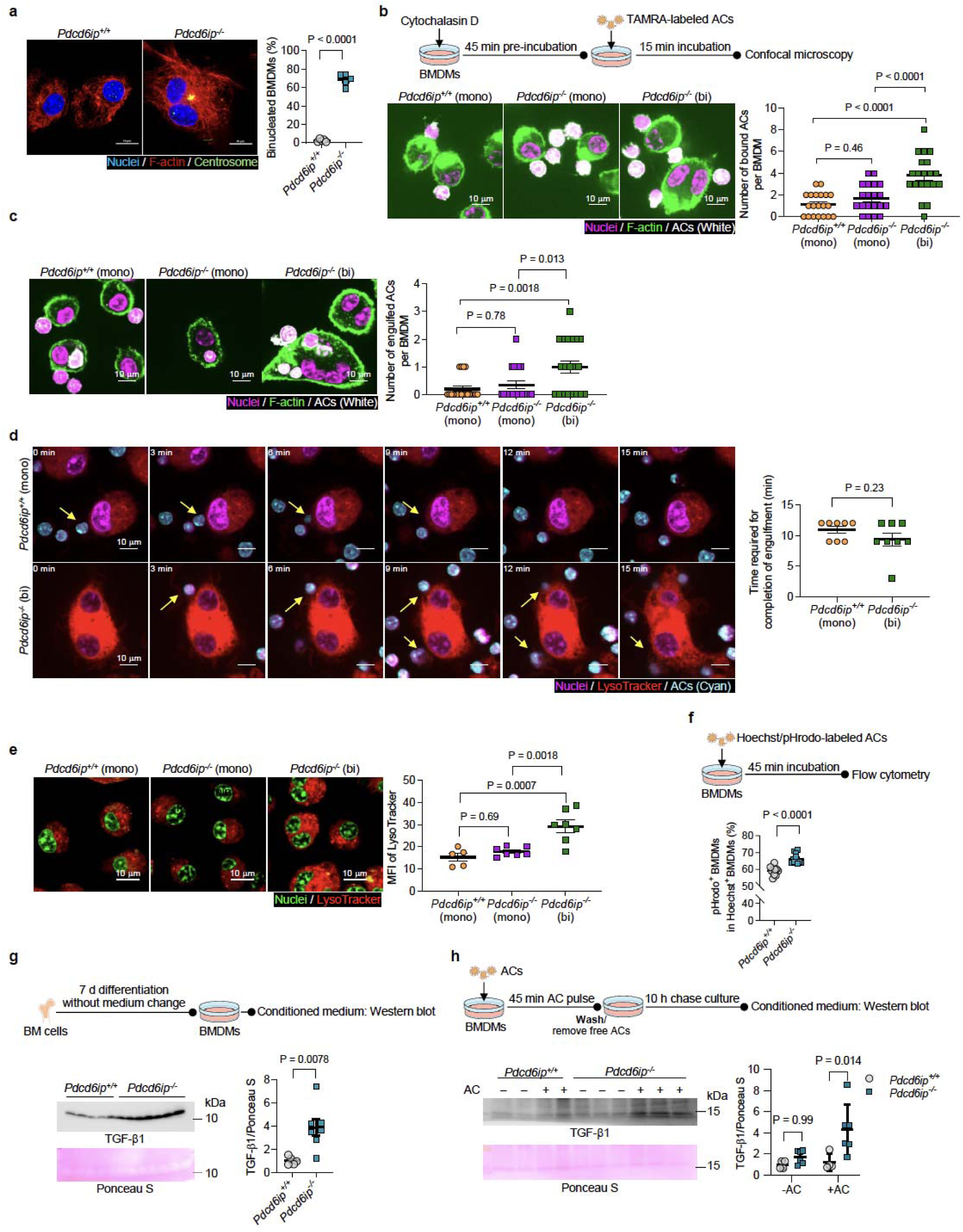
*Pdcd6ip* deletion induces the formation of binucleated macrophages with enhanced apoptotic cell binding, engulfment, lysosomal acidification, and resolution response. **a**, Representative confocal images and quantification of binucleated *Pdcd6ip*^+/+^ and *Pdcd6ip*^−/−^ BMDMs. *In vitro* cultured BMDMs were fixed and stained for nuclei (Hoechst), F-actin (Phalloidin), and centrosomes (Pericentrin). *n* = 5 *Pdcd6ip^+/+^* and 5 *Pdcd6ip*^−/−^ mice. **b,** Quantification of AC binding. BMDMs were pretreated with cytochalasin D, an actin polymerization inhibitor, to block internalization, followed by incubation with TAMRA-labeled ACs for 15 min. Cells were then washed to remove unbound ACs, fixed, stained for nuclei (DRAQ5) and F-actin (Phalloidin), and imaged by confocal microscopy. *Pdcd6ip*^−/−^ binucleated (bi) BMDMs showed increased AC binding. Twenty cells per group from 3 *Pdcd6ip*^+/+^ and 3 *Pdcd6ip*^−/−^ mice were analyzed. **c,** Quantification of AC engulfment by BMDMs with a 45 min incubation showed increased multi-corpse engulfment in *Pdcd6ip*^−/−^ (bi) BMDMs. Cells were stained and imaged as described in **b**. Twenty cells per group from 3 *Pdcd6ip*^+/+^ and 3 *Pdcd6ip*^−/−^ mice were analyzed. **d,** Time-lapse live-cell imaging showed no significant difference in the time from AC binding to completion of engulfment between *Pdcd6ip*^+/+^ and *Pdcd6ip*^−/−^ BMDMs. Representative time-lapse frames showing the transition from initial apoptotic cell binding to complete engulfment at the indicated time points. Yellow arrows mark the apoptotic cell followed throughout the imaging sequence. ACs were labeled by Hoechst and nuclei were stained by DRAQ5. Eight cells per group from *n* = 3 *Pdcd6ip*^+/+^ and 3 *Pdcd6ip*^−/−^ mice. **e,** Representative live-cell images and quantification of LysoTracker staining showed increased lysosomal signal in *Pdcd6ip*^−/−^ (bi) BMDMs. Twenty to thirty cells were quantified per mouse, and the mean value for each mouse was used for analysis. *n* = 5 *Pdcd6ip*^+/+^ and 7 *Pdcd6ip*^−/−^ mice. **f,** Flow cytometry analysis of efferocytosis of Hoechst and pHrodo dual-labeled ACs showed increased acidification of engulfed AC cargo in *Pdcd6ip*^−/−^ BMDMs. ACs were dual-labeled with Hoechst (pH-insensitive uptake marker) and pHrodo probe (pH-sensitive acidification maker), and co-incubated with BMDMs for 45 min. Hoechst^+^ BMDMs were quantified as AC-engulfing cells, whereas pHrodo positivity among Hoechst^+^ BMDMs was used to assess acidification of engulfed ACs. *n* = 10 *Pdcd6ip*^+/+^ and 11 *Pdcd6ip*^−/−^ mice. **g and h,** Western blot analysis of secreted TGF-β1 in conditioned medium from BMDMs cultured for seven days without media change (**g**) or incubated with ACs for 45 min, followed by AC removal and an additional 10 h culture period (**h**). *Pdcd6ip*^−/−^ BMDMs showed increased TGF-β1 secretion under both conditions. *n* = 6 *Pdcd6ip*^+/+^ and 8 *Pdcd6ip*^−/−^ mice in panel **g**, and *n* = 4 *Pdcd6ip*^+/+^ and 6 *Pdcd6ip*^−/−^ mice in panel **h**. Scale bar = 10 μm in all representative images. Data are shown as mean ± SEM. Statistical significance was determined using a two-tailed Student’s t-test for **a**, **d**, **f**, and **g**; one-way ANOVA with Tukey’s post hoc test was used for **b**, **c**, **e**; and two-way ANOVA with Bonferroni’s post hoc test was used for **h**.

We next asked whether the increased efferocytosis observed in *Pdcd6ip^−/−^*BMDMs is linked specifically to binucleation. To distinguish AC binding from engulfment, BMDMs were pretreated with cytochalasin D to prevent actin-dependent internalization and then incubated with ACs. Under these conditions, binucleated *Pdcd6ip^−/−^* macrophages bound more ACs per macrophage than *Pdcd6ip^+/+^* macrophages, whereas mononuclear *Pdcd6ip^−/−^* macrophages showed no difference compared with *Pdcd6ip^+/+^* controls (**Fig. 2b**). Imaging-based analyses further assessed the capacity of individual macrophages to engulf multiple ACs. Binucleated *Pdcd6ip^−/−^*macrophages more frequently contained multiple ACs per macrophage, whereas mononuclear *Pdcd6ip^−/−^* macrophages did not differ substantially from *Pdcd6ip^+/+^* macrophages (**Fig. 2c**). Thus, the enhanced efferocytic phenotype caused by *Pdcd6ip* deletion was driven by the binucleated macrophage population. Importantly, when the time from AC binding to completion of engulfment was quantified, binucleated *Pdcd6ip^−/−^* macrophages did not engulf individual ACs faster than *Pdcd6ip^+/+^* macrophages (**Fig. 2d**). These findings suggest that binucleated *Pdcd6ip^−/−^* macrophages enhance efferocytosis primarily by increasing AC binding and multi-cargo engulfment capacity rather than by accelerating the engulfment of each individual target.

The altered side-scatter profile of *Pdcd6ip^−/−^* macrophages suggested increased intracellular granularity, which we hypothesized could reflect expanded lysosomal content. Consistent with this idea, binucleated *Pdcd6ip^−/−^*macrophages showed increased LysoTracker signal compared with mononuclear *Pdcd6ip^−/−^*and *Pdcd6ip^+/+^* macrophages (**Fig. 2e**). Because efficient efferocytosis requires not only AC engulfment but also phagolysosome maturation and cargo degradation, we next quantified lysosomal acidification of engulfed cargo using ACs dual-labeled with Hoechst, a pH-insensitive dye, and pHrodo, a pH-sensitive dye. Hoechst positivity marked macrophages that had engulfed ACs, whereas dual Hoechst and pHrodo positivity marked engulfed cargo that had entered an acidic compartment^31^. *Pdcd6ip^−/−^* macrophages showed increased pHrodo positivity among Hoechst-positive macrophages, indicating enhanced acidification normalized to engulfment (**Fig. 2f**).

Finally, we assessed whether enhanced efferocytosis is accompanied by a resolution-associated secretory response. In extended BMDM cultures maintained for seven days without media change, *Pdcd6ip^−/−^*macrophages accumulated higher TGF-β1 in conditioned media than *Pdcd6ip^+/+^*macrophages, likely reflecting increased clearance of dying cells generated during culture (**Fig. 2g**). In a more direct AC feeding assay, *Pdcd6ip^−/−^* macrophages did not show increased TGF-β1 compared with *Pdcd6ip^+/+^* macrophages under basal short-term conditions but produced more TGF-β1 after AC exposure (**Fig. 2h**). Because TGF-β1 is a canonical pro-reparative and anti-inflammatory mediator induced by efferocytosis and contributing to resolution^32^, these data indicate that *Pdcd6ip^−/−^* macrophages couple enhanced AC uptake and degradation to an enhanced resolution response.

To distinguish whether acute modulation of PDCD6IP protein abundance directly alters efferocytosis independent of binucleation, we performed PDCD6IP overexpression experiments. PDCD6IP overexpression did not alter engulfment, but modestly reduced acidification (**Supplementary Fig. 2**), supporting the model in which *Pdcd6ip* loss enhances efferocytosis primarily by inducing binucleation and associated macrophage-state reprogramming rather than through an acute, direct effect of PDCD6IP abundance on the efferocytic machinery.

### Single-cell RNA sequencing reveals a distinct transcriptomic signature associated with binucleated *Pdcd6ip^−/−^* macrophages

The imaging data suggested that binucleated *Pdcd6ip^−/−^*macrophages are functionally distinct, prompting us to ask whether this phenotype reflects only increased cell size and nuclear content or is associated with a distinct transcriptomic profile. We therefore performed scRNA-seq of sorted BMDM populations enriched for mononuclear *Pdcd6ip^+/+^*BMDMs, mononuclear *Pdcd6ip^−/−^* BMDMs, and binucleated *Pdcd6ip^−/−^*BMDMs. Samples were indexed by hashing before single-cell capture and sequencing (**Fig. 3a**, schematic of the experimental design). Because conventional fluorescence-activated cell sorting provided better BMDM viability under our experimental conditions, we used the live-cell DNA dye DyeCycle Green together with forward scatter to sort populations differing in nuclear content and cell size. DyeCycle Green^hi^/FSC-A^hi^ and DyeCycle-Green^low^/FSC-A^low^ BMDMs were collected as binucleated cell-enriched and mononuclear cell-enriched populations, respectively. *Pdcd6ip^−/−^* BMDMs contained a higher proportion of DyeCycle Green^hi^/FSC-A^hi^ cells than *Pdcd6ip^+/+^*BMDMs, consistent with binucleated cell enrichment in this sorted population (**Fig. 3b**). Because DNA content cannot fully distinguish replicating mononuclear cells from truly binucleated cells, this approach provided enrichment rather than absolute purification. The success of enrichment is confirmed by microscopy imaging of the sorted populations (**Fig. 3b**).

**Fig. 3.**
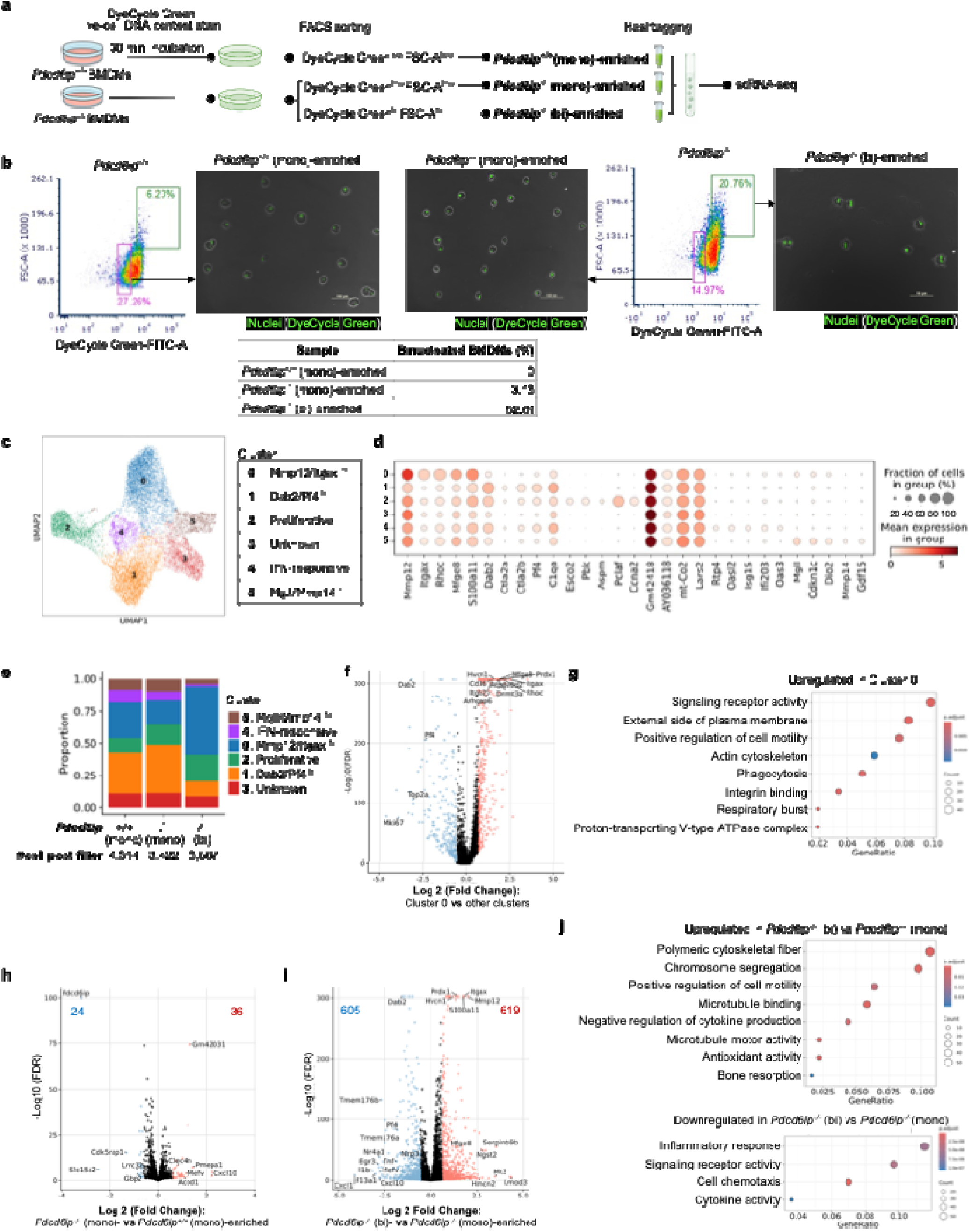
Single-cell transcriptomic profiling reveals a distinct transcriptomic signature associated with binucleated *Pdcd6ip^−/−^* macrophages. **a**, Schematic of the single-cell RNA sequencing (scRNA-seq) experimental design. BMDMs were generated from *Pdcd6ip^+/+^* and *Pdcd6ip^−/−^* mice, stained with DyeCycle Green to assess DNA content, and sorted using DyeCycle Green intensity together with forward scatter to enrich mononuclear (mono) and binucleated (bi) BMDM fractions. Cells were indexed by sample hashing before single-cell capture and sequencing. **b,** Flow cytometry sorting strategy of DyeCycle Green^hi^/FSC-A^hi^ cells and DyeCycle Green^low^/FSC-A^low^ cells. DyeCycle Green^hi^/FSC-A^hi^ cells were sorted as binucleated cell-enriched BMDM population, whereas DyeCycle Green^low^/FSC-A^low^ cells were sorted as mononuclear cell-enriched BMDM population. The enrichment was confirmed by microscopy imaging of sorted cells. *Pdcd6ip^−/−^* BMDMs contained a higher fraction of DyeCycle Green^hi^/FSC-A^hi^ cells than *Pdcd6ip^+/+^* BMDMs. Each genotype was represented by a pooled sample generated from three mice and profiled in two 10x Chromium runs. **c,** Uniform manifold approximation and projection (UMAP) visualization of scRNA-seq profiles from sorted BMDM populations. **d,** Dot plot showing expression of representative marker genes across annotated BMDM clusters. Complete differential expression results are provided in **Supplementary Table 1**. **e,** Quantification of cluster proportions across sorted populations showing preferential enrichment of Cluster 0 in the binucleated cell-enriched *Pdcd6ip^−/−^* BMDM fraction. Complete analysis is provided in **Supplementary Table 2**. **f,** Volcano plot showing differentially expressed genes in Cluster 0 compared with all other clusters, including genes known to regulate efferocytosis. **g,** Gene Ontology analysis of genes upregulated in Cluster 0. Complete Gene Ontology results are provided in in **Supplementary Table 3**. **h,** Pseudobulk differential expression analysis comparing mononuclear cell-enriched *Pdcd6ip^+/+^*and *Pdcd6ip^−/−^* BMDMs, showing overall similar transcriptomic profiles. Complete differential expression results are provided in **Supplementary Table 4**. **i,** Pseudobulk differential expression analysis comparing binucleated cell-enriched *Pdcd6ip^−/−^* and mononuclear cell-enriched *Pdcd6ip^−/−^* BMDMs, showing a distinct transcriptomic profile in the binucleated cell-enriched fration. Complete differential expression results are provided in **Supplementary Table 4**. **j,** Gene Ontology analysis of differentially expressed genes between binucleated cell-enriched and mononuclear cell-enriched *Pdcd6ip^−/−^*BMDMs. Complete Gene Ontology results are provided in **Supplementary Table 5**.

Unsupervised clustering analysis identified six BMDM clusters (**Fig. 3c**), showing overall transcriptomic similarity among several clusters, with a distinct proliferative subpopulation, consistent with previous BMDM scRNA-seq studies^33^. Expression of representative marker genes used to annotate the clusters is shown in **Fig. 3d** and **Supplementary Table 1**. In addition, we identified a distinct subpopulation, Cluster 0, that was preferentially enriched in the binucleated cell-enriched *Pdcd6ip^−/−^*BMDM population (**Fig. 3e** and **Supplementary Table 2**). Cluster 0 subpopulation showed higher expression of multiple genes known to regulate integrin signaling and surface recognition (*Itgax*, *Cd36*, *Itgb2*, *Mfge8*), actin remodeling (*Rhoc*, *Arhgap6*), lysosomal acidification (*Hvcn1*, *Atp6v0d2*), and anti-oxidative and resolution response (*Prdx1*, *Dnmt3a*) (**Fig. 3f**). Gene ontology analysis of upregulated genes in Cluster 0 cells indeed showed enrichment for genes annotated to gene ontology (GO) terms, including phagocytosis,integrin signaling, actin cytoskeleton, and respiratory burst (**Fig. 3g** and **Supplementary Table 3**).

To define this transcriptomic signature more robustly, we performed pseudobulk differential expression analyses across the sorted populations. Mononuclear cell-enriched *Pdcd6ip^+/+^* and *Pdcd6ip^−/−^*BMDMs were broadly similar at the transcriptomic level, with few differentially expressed (DE) genes, aside from the expected reduction of *Pdcd6ip* expression in *Pdcd6ip^−/−^* BMDMs (**Fig. 3h** and **Supplementary Table 4**). In contrast, binucleated cell-enriched *Pdcd6ip^−/−^* BMDMs showed a distinct transcriptomic profile compared with mononuclear cell-enriched *Pdcd6ip^−/−^*BMDMs, with numerous upregulated and downregulated DE genes (**Fig. 3i** and **Supplementary Table 4**). Upregulated genes include multiple Cluster 0 markers (**Fig. 3f**), whereas downregulated genes include Cluster 1 markers (*Dab2*, *Pf4*) (**Fig. 3d**), indicating that differences in cluster composition contribute to the transcriptomic divergence between the sorted populations.

Gene ontology analysis of genes upregulated in binucleated cell-enriched *Pdcd6ip^−/−^* BMDMs showed enrichment for genes annotated to chromosome segregation, positive regulation of cell motility, negative regulation of cytokine production, bone resorption, polymeric cytoskeletal fiber, microtubule binding, microtubule motor activity, antioxidant activity (**Fig. 3j** and **Supplementary Table 5**). Gene ontology analysis of downregulated genes in binucleated cell-enriched *Pdcd6ip^−/−^* BMDMs showed enrichment for genes annotated to cell chemotaxis, inflammatory response, signaling receptor activity, and cytokine activity (**Fig. 3j** and **Supplementary Table 5**). Together, these findings indicate that *Pdcd6ip* loss is associated not simply with increase macrophage size or nuclear content, but with a distinct binucleation-associated transcriptomic state, characterized by altered cytoskeletal and motility programs, enhanced antioxidant capacity, and reduced inflammatory signaling.

### *Pdcd6ip* deletion enhances *in vivo* apoptotic cell clearance and protects mice from autoimmune responses to chronic dead cell burden

We next asked whether *Pdcd6ip* deletion increases AC clearance *in vivo*. Fluorescently labeled ACs were injected into *Pdcd6ip^+/+^*and *Pdcd6ip^−/−^* mice, and splenic macrophage uptake was quantified 24 and 48 h later by measuring TAMRA-positive CD45^+^CD11b^+^F4/80^+^ splenic macrophages (**Supplementary Fig. 3a**). At 24 h after injection, *Pdcd6ip^−/−^*mice had a higher fraction of TAMRA-positive splenic macrophages than *Pdcd6ip^+/+^*mice, indicating enhanced *in vivo* uptake of ACs (**Supplementary Fig. 3b**). By 48 h, this difference was no longer evident, consistent with subsequent degradation of engulfed cargo and clearance of the fluorescent signal (**Supplementary Fig. 3c**).

To determine whether enhanced clearance is protective under conditions of sustained dead cell burden, we used a repeated AC injection model (**Fig. 4a**). In our prior work, repeated delivery of ACs at a dose exceeding endogenous clearance capacity induces mild autoimmunity in C57BL/6 mice, characterized by antinuclear antibody (ANA) generation and immune-complex deposition in kidney glomeruli^21^. Using this model, *Pdcd6ip^−/−^* mice showed reduced circulating ANA levels compared with *Pdcd6ip^+/+^* mice, whereas anti-dsDNA levels remained low and not significantly different, consistent with the limited anti-dsDNA induction in this model^21^ (**Fig. 4b,c**). Histological analysis revealed a more pronounced protective effect of *Pdcd6ip^−/−^* in the kidney. Repeated AC injection increased glomerular immune-complex deposition, but *Pdcd6ip^−/−^* mice had significantly lower glomerular IgG and C1q deposition than *Pdcd6ip^+/+^* mice (**Fig. 4d,e**). These findings indicate that *Pdcd6ip* deletion enhances *in vivo* AC clearance and protects mice from autoimmune responses caused by chronic dead cell burden.

**Fig. 4.**
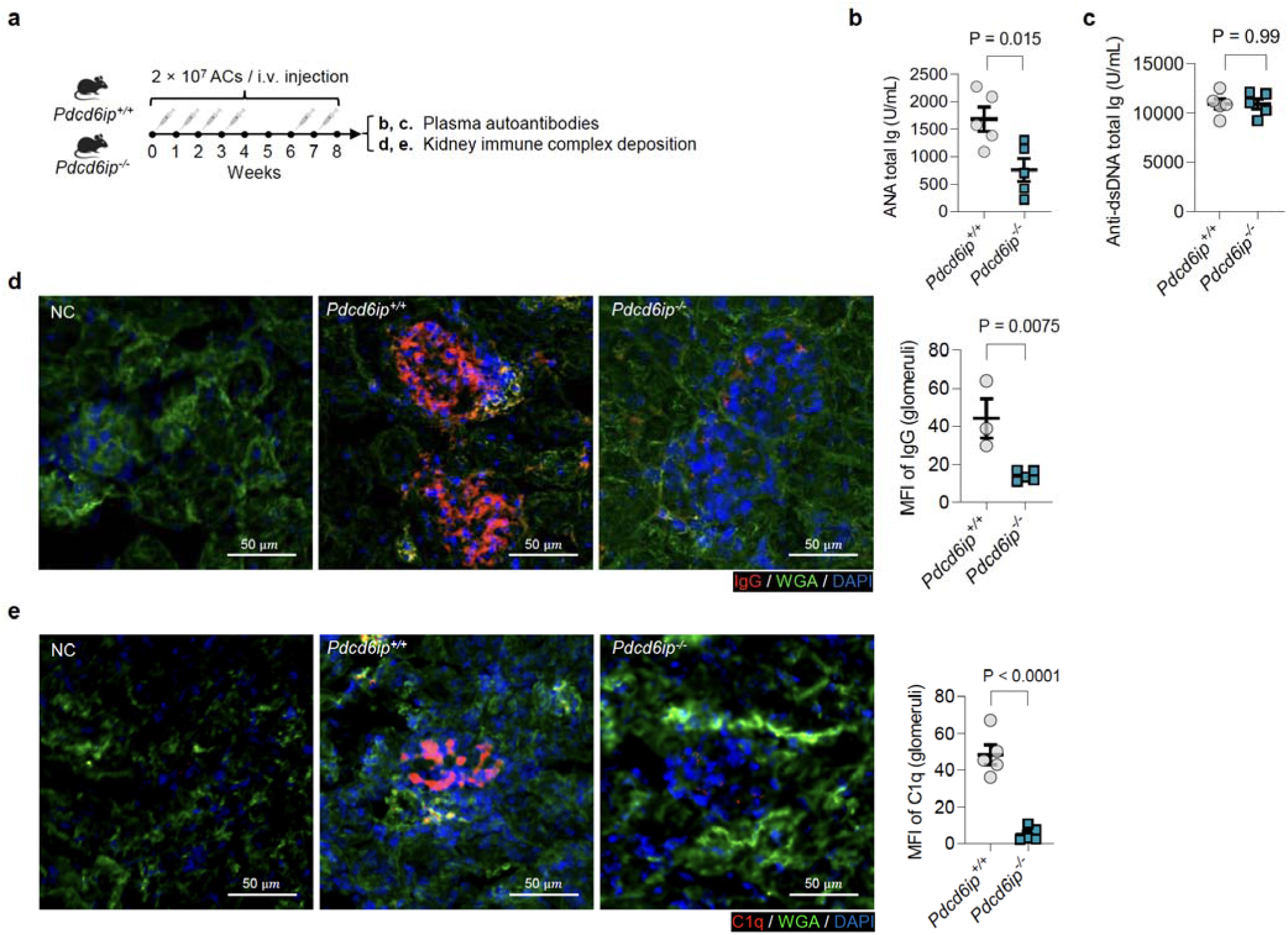
*Pdcd6ip* deletion protects mice from autoimmune responses induced by chronic apoptotic cell burden. **a**, Schematic of the experimental design. A repeated AC injection model was used to induce sustained dead cell burden and autoimmune responses. ACs were injected intravenously via the tail vein into *Pdcd6ip*^+/+^ and *Pdcd6ip*^−/−^ mice according to the indicated timeline. Plasma samples were collected to measure anti-nuclear antibodies (ANA) and anti-dsDNA antibodies. Kidneys were dissected and sectioned for immunofluorescence staining of immune complex deposition. **b, c**, Quantification of ANA total Ig and anti-dsDNA total Ig in plasma by ELISA. *Pdcd6ip*^−/−^ mice showed reduced ANA levels compared with *Pdcd6ip*^+/+^ mice, whereas anti-dsDNA levels remained low and were not significantly different between groups. *n* = 5 *Pdcd6ip*^+/+^ and 5 *Pdcd6ip*^−/−^ mice. **d, e**, Immunofluorescence staining of kidney sections using anti-IgG or anti-C1q antibodies, co-stained with wheat germ agglutinin (WGA) to outline glomeruli and DAPI to label nuclei. *Pdcd6ip*^+/+^mice showed reduced glomerular IgG and C1q deposition compared with *Pdcd6ip*^+/+^ mice. *n* = 3 *Pdcd6ip*^+/+^ and 5 *Pdcd6ip*^−/−^ mice for **d**; *n* = 5 *Pdcd6ip*^+/+^ and 5 *Pdcd6ip*^−/−^ mice for **e**. Data are shown as mean ± SEM. Statistical significance was determined using a two-tailed Student’s *t*-test.

### Hematopoietic *Pdcd6ip* deletion stabilizes atherosclerotic plaques without altering systemic parameters

Atherosclerosis provides an important disease context in which defective macrophage efferocytosis has direct pathological consequences^4^. Human plaque studies showed that clearance of dying cells and associated debris by macrophages is impaired in advanced atherosclerotic lesions^34^, and mouse studies have established that disruption of efferocytosis pathways promotes apoptotic cell accumulation, necrotic core expansion, and features of plaque instability^35–37^. Conversely, enhancing macrophage efferocytosis can limit necrotic core formation and improve features associated with plaque stability^38^. These observations have motivated therapeutic strategies that seek not only to reduce lipid burden but also to restore defective clearance and resolution within the plaque microenvironment^4^. We therefore next tested whether the enhanced efferocytic capacity caused by hematopoietic *Pdcd6ip* deletion protects against atherosclerosis. *Ldlr^−/−^* recipient mice were lethally irradiated and reconstituted with bone marrow from *Pdcd6ip^+/+^*or *Pdcd6ip^−/−^* donor mice, followed by Western diet feeding for 21-23 weeks to induce advanced atherosclerotic lesions (**Fig. 5a**, schematic of the experimental design). Successful hematopoietic reconstitution with *Pdcd6ip^−/−^* bone marrow was confirmed by reduced PDCD6IP protein abundance in BMDMs from recipient mice (**Fig. 5b**).

**Fig. 5.**
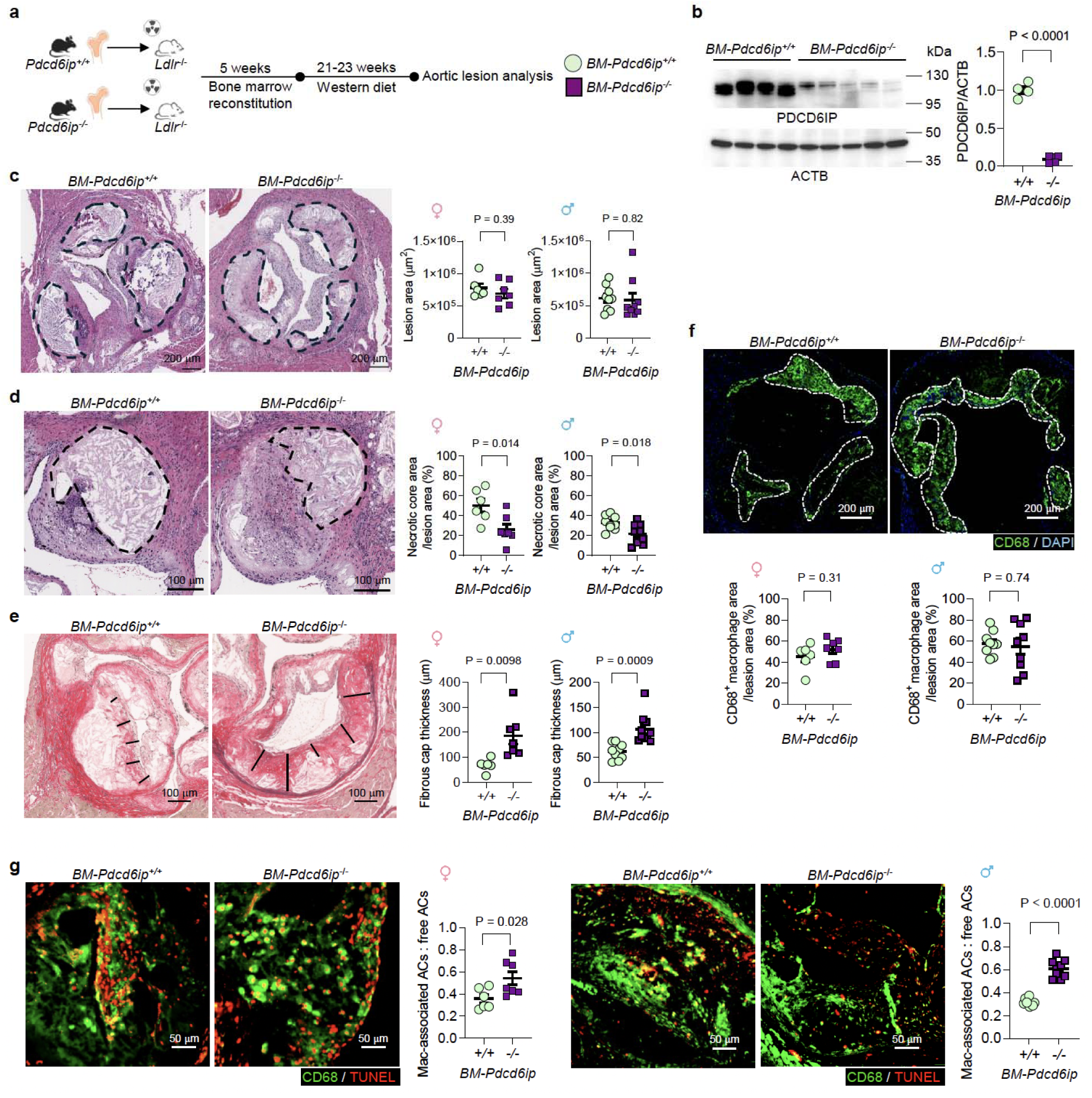
Hematopoietic *Pdcd6ip* deletion promotes plaque stability and enhances lesional efferocytosis in *Ldlr*^−/−^ mice. **a**, Schematic of the study design. *Ldlr*^−/−^ recipient mice were lethally irradiated and transplanted with bone marrow from *Pdcd6ip^+/+^*or *Pdcd6ip^−/−^* donor mice. After 5 weeks of hematopoietic reconstitution, mice were fed a Western diet for 21-23 weeks. Throughout this figure, *BM-Pdcd6ip^+/+^* and *BM-Pdcd6ip^−/−^*indicate *Ldlr^−/−^* recipient mice reconstituted with *Pdcd6ip^+/+^*or *Pdcd6ip^−/−^* bone marrow, respectively. **b,** Western blot analysis confirming reduced PDCD6IP protein in BMDMs generated from *BM-Pdcd6ip^−/−^*recipient mice, compared with *BM-Pdcd6ip^+/+^* recipient mice. *n* = 4 *BM-Pdcd6ip^+/+^* and 5 *BM-Pdcd6ip^−/−^*female mice. **c,** Quantification of lesion area by H&E staining. Hematopoietic *Pdcd6ip* deletion did not significantly alter the total lesion area. Female: *n* = 6 *BM-Pdcd6ip^+/+^* and 7 *BM-Pdcd6ip^−/−^*mice; male: *n* = 9 *BM-Pdcd6ip^+/+^* and 9 *BM-Pdcd6ip^−/−^*mice. Scale bar = 200 μm. **d,** Quantification of necrotic core area by H&E staining, normalized to lesion area, showing reduced necrotic core size in *BM-Pdcd6ip^−/−^* recipient mice. Female: *n* = 6 *BM-Pdcd6ip^+/+^* and 7 *BM-Pdcd6ip^−/−^*mice; male: *n* = 9 *BM-Pdcd6ip^+/+^* and 9 *BM-Pdcd6ip^−/−^*mice. Scale bar = 100 μm. **e,** Quantification of fibrous cap thickness by Picrosirius red staining showing increased fibrous cap thickness in *BM-Pdcd6ip^−/−^* recipient mice. Female: *n* = 6 *BM-Pdcd6ip^+/+^* and 7 *BM-Pdcd6ip^−/−^* mice; male: *n* = 9 *BM-Pdcd6ip^+/+^* and 10 *BM-Pdcd6ip^−/−^*mice. Scale bar = 100 μm. **f,** Quantification of CD68^+^ plaque macrophage area by immunofluorescence staining showing no significant difference between *BM-Pdcd6ip^+/+^* and *BM-Pdcd6ip^−/−^*recipient mice. CD68 (green) and DAPI (blue). CD68^+^ macrophage area was normalized to lesion area. Female: *n* = 6 *BM-Pdcd6ip^+/+^*and 7 *BM-Pdcd6ip^−/−^* mice; male: *n* = 9 *BM-Pdcd6ip^+/+^*and 9 *BM-Pdcd6ip^−/−^* mice. Scale bar = 200 μm. **g**, *In situ* efferocytosis analysis in atherosclerotic plaques. Apoptotic cells or fragments were detected by TUNEL staining, and macrophages were identified by CD68 staining. The efferocytosis index was calculated as the ratio of macrophage-associated to free TUNEL+ apoptotic cells or fragments. Hematopoietic *Pdcd6ip* deletion increased the *in situ* efferocytosis index in both female and male recipient mice. Female: *n* = 6 *BM-Pdcd6ip^+/+^* and *n* = 7 *BM-Pdcd6ip^−/−^* mice; male: *n* = 9 *BM-Pdcd6ip^+/+^*and 9 *BM-Pdcd6ip^−/−^* mice. Scale bar = 50 μm. Data are shown as mean ± SEM. Statistical significance was determined using a two-tailed Student’s t-test.

Atherosclerotic lesion analysis showed that hematopoietic *Pdcd6ip* deletion did not significantly change total lesion area in either female or male recipients (**Fig. 5c**). However, *Pdcd6ip^−/−^*bone marrow recipients displayed smaller necrotic cores and increased fibrous cap thickness compared with *Pdcd6ip^+/+^* bone marrow recipients (**Fig. 5d,e**). These features indicate improved plaque stability despite unchanged overall lesion size. Macrophage area within lesions was not substantially altered (**Fig. 5f**), suggesting that the protective phenotype was not simply caused by reduced macrophage accumulation. We identified apoptotic/dead cells in plaque sections by terminal deoxynucleotidyl transferase dUTP nick-end labeling (TUNEL), which detects fragmented DNA, and quantified *in situ* efferocytosis as the ratio of macrophage-associated to free TUNEL^+^ ACs or AC fragments. ACs in direct contact with macrophages were interpreted as macrophage-associated and likely undergoing clearance, whereas ACs not associated with macrophages were interpreted as free apoptotic bodies. Hematopoietic *Pdcd6ip* deletion increased this *in situ* efferocytosis index in both female and male recipient mice (**Fig. 5g**). Consistent with enhanced efferocytosis and inflammation resolution, *Pdcd6ip^−/−^* bone marrow recipients also showed increased plaque TGF-β1, a resolution-associated anti-inflammatory mediator linked to fibrous cap stability, and reduced IL-1β, a pro-inflammatory cytokine implicated in plaque inflammation (**Supplementary Fig. 4**). Together, these data suggest that hematopoietic *Pdcd6ip* deletion stabilizes plaques by enhancing lesional efferocytosis and shifting the local inflammatory milieu toward resolution.

The atheroprotective phenotype occurred without major changes in systemic parameters. Body weight, plasma cholesterol levels, and Complete Blood Count with differential were comparable between *Pdcd6ip^+/+^* and *Pdcd6ip^−/−^* bone marrow recipients (**Supplementary Fig. 5**). Thus, hematopoietic *Pdcd6ip* deletion improved plaque stability features independently of changes in systemic lipid exposure or circulating leukocyte abundance.

### Binucleated *Pdcd6ip^−/−^* macrophages are generated *in vivo* and retain enhanced efferocytic capacity *ex vivo*

The atherosclerosis data raised the question of whether binucleated *Pdcd6ip^−/−^* macrophages are present *in vivo* and retain the pro-efferocytic phenotype observed in cultured BMDMs. Because atherosclerotic plaques yield limited macrophage numbers and two-dimensional histological sections cannot reliably determine whether individual macrophages are binucleated, we focused on splenic macrophages from bone marrow transplant recipient mice as an accessible *in vivo* macrophage population (**Fig. 6a**, schematic of the experimental design).

**Fig. 6.**
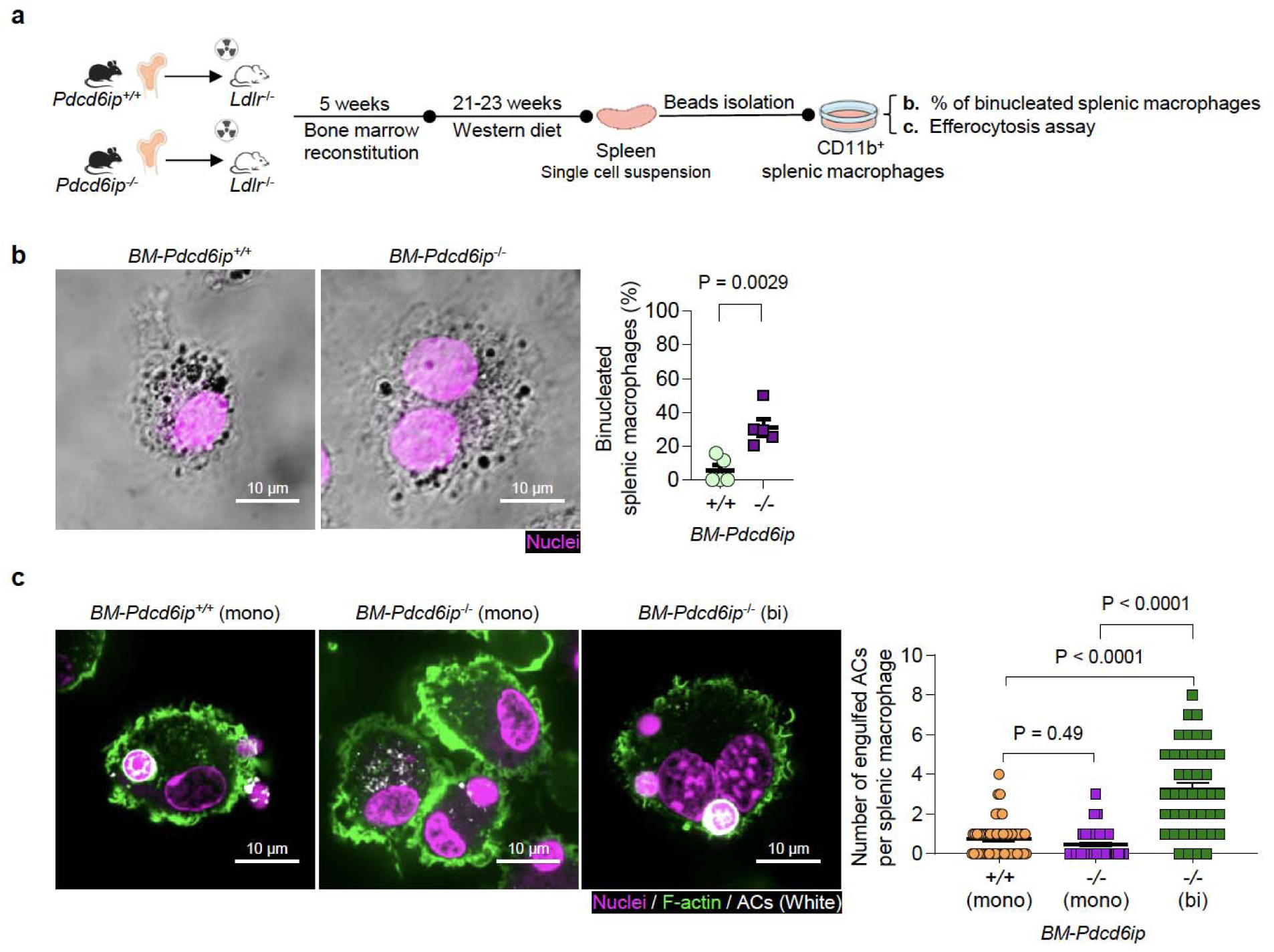
Binucleated *Pdcd6ip^−/−^*macrophages arise *in vivo* and retain enhanced efferocytic capacity *ex vivo*. **a**, Schematic of the experimental design used to isolate and analyze splenic macrophages from bone marrow transplant recipient mice, as in **Fig. 4**. Splenic macrophages were isolated from *Pdcd6ip^+/+^*and *Pdcd6ip^−/−^* bone marrow recipient mice by bead-based enrichment of CD11b^+^ cells, plated for culture of 6-8 h to allow adherence, followed by nuclear staining and live-cell imaging by confocal microscopy or for *ex vivo* efferocytosis assay. **b,** Representative live-cell images showing binucleated macrophages among splenic macrophages isolated from *Pdcd6ip^−/−^* bone marrow recipient mice (*BM-Pdcd6ip^−/−^*). Scale bar = 10 μm. *n* = 5 *BM-Pdcd6ip^+/+^*and 5 *BM-Pdcd6ip^−/−^* male mice. **c,** *Ex vivo* efferocytosis assay of splenic macrophages. CD11b^+^ splenic macrophages were incubated with TAMRA-labeled ACs for 45 min. Cells were then washed to remove unbound ACs, fixed, stained for nuclei (DRAQ5) and F-actin (Phalloidin), and imaged by confocal microscopy. Binucleated *Pdcd6ip^−/−^* splenic macrophages engulfed more ACs per macrophage than *Pdcd6ip^+/+^* macrophages. Mononuclear *Pdcd6ip^−/−^* splenic macrophages were comparable to *Pdcd6ip^+/+^* macrophages. *n* = 58 mononuclear cells from 5 *Pdcd6ip^+/+^*, 58 mononuclear cells from 5 *BM-Pdcd6ip^−/−^*, and 46 binucleated cells from 5 *BM-BM-Pdcd6ip^−/−^*male mice. Scale bar = 10 μm. Data are shown as mean ± SEM. Statistical significance was determined using a two-tailed Student’s t-test in panel **b** or one-way ANOVA followed by Tukey’s post hoc test in panel **c**.

Splenic macrophages were isolated from *Pdcd6ip^+/+^* and *Pdcd6ip^−/−^* bone marrow recipient mice using bead-based enrichment, plated to allow adherence, and analyzed by microscopy. Binucleated macrophages were readily detected among splenic macrophages from *Pdcd6ip^−/−^*bone marrow recipients (**Fig. 6b**). *Ex vivo* efferocytosis assays showed that binucleated *Pdcd6ip^−/−^* splenic macrophages engulfed more ACs per macrophage than *Pdcd6ip^+/+^* macrophages, including cells containing multiple apoptotic targets and, in some cases, five to eight ACs per macrophage (**Fig. 6c**). By contrast, mononuclear *Pdcd6ip^−/−^* splenic macrophages were comparable to *Pdcd6ip^+/+^* macrophages (**Fig. 6c**). These findings demonstrated that *Pdcd6ip* deletion generates binucleated macrophages *in vivo* that possess enhanced efferocytic capacity. Bulk RNA-seq of whole spleen followed by differential expression analysis revealed minimal transcriptomic differences between *Pdcd6ip^+/+^*and *Pdcd6ip^−/−^* bone marrow recipient mice with only 4 DE genes (an absolute fold change ≥1.5, FDR <0.05, **Supplementary Fig. 6** and **Supplementary Table 6**), suggesting that hematopoietic *Pdcd6ip* deletion does not substantially alter the overall splenic transcriptomic state.

### *PDCD6IP* perturbation generates binucleated, pro-efferocytic human macrophage-like cells

Finally, we asked whether *PDCD6IP* perturbation could be used to generate binucleated, pro-efferocytic human macrophage-like cells. As a proof-of-principle experiment, THP-1 monocytes were transduced with lentiviral vectors encoding Cas9 and *PDCD6IP*-targeting sgRNA, followed by puromycin selection to enrich edited cells (**Supplementary Fig. 7a**, schematic of the experimental design). This strategy enriched a binucleated THP-1 monocyte population (**Supplementary Fig. 7b**) that could subsequently be differentiated into binucleated macrophage-like cells (**Supplementary Fig. 7c**).

Although THP-1-derived macrophage-like cells possess low basal efferocytic capacity^20^, *PDCD6IP*-targeted cells showed increased AC engulfment compared with non-targeting sgRNA controls (**Supplementary Fig. 7d**). Consistent with the murine BMDM phenotype, binucleated *PDCD6IP*-deleted THP-1 macrophage-like cells showed increased numbers of engulfed ACs per macrophage (**Supplementary Fig. 7d**). They also showed increased TGF-β1 expression (**Supplementary Fig. 7e**). These data suggest that *PDCD6IP* perturbation can generate binucleated human macrophage-like cells with enhanced engulfment and resolution-associated responses, supporting further development of strategies to engineer pro-efferocytic macrophages.

## DISCUSSION

Our study demonstrates that *Pdcd6ip* deletion reprograms macrophages into a binucleated, highly efferocytic state, driving coordinated improvements across multiple stages of apoptotic cell clearance and downstream resolution. *In vivo*, this genetic perturbation reduces immune-complex deposition in a chronic apoptotic cell burden model and stabilizes advanced atherosclerotic plaques. By acquiring binucleation via cytokinesis arrest, a mechanism distinct from classical, fusion-mediated MGC formation^10–12^, macrophages achieve a functional state uniquely specialized for enhanced efferocytosis. Together, these findings reveal a previously unrecognized link between cytokinesis regulation, macrophage cell architecture, efferocytic capacity, and disease protection.

Mechanistically, systemic loss of PDCD6IP is linked to developmental abnormalities such as microcephaly^39,40^. Homozygous knockout mice are recovered below the expected Mendelian frequency at birth despite normal Mendelian ratios at embryonic day 12.5, suggesting partially penetrant lethality later in development^39^. Our findings show that macrophages are relatively permissive to PDCD6IP loss and incomplete cytokinesis. Rather than undergoing cell death, macrophages functionally tolerate binucleation and acquired enhanced efferocytic capacity. These results suggest that the functional consequences of cytokinetic failure are likely dependent on cellular context.

These findings also fit within the broader, emerging biology of polyploid cells. Polyploidy is a recognized feature in hepatocytes, cardiomyocytes, and osteoclasts^18,41–44^, yet the physiological functions of many multinucleated or polyploid states remain incompletely defined. In some contexts, polyploidy has been linked to stress tolerance, wound repair, metabolic adaptation, and disease prognosis^41,44–49^, or acquisition of phagocytic capacity^13–18,50^. Our study adds macrophage efferocytosis to this conceptual framework, suggesting that cytokinesis arrest can be repurposed by macrophages to meet specialized functional demands for increased dying cell clearance, a mechanism with important implications for autoimmunity, chronic inflammation, aging, and atherosclerotic progression.

Translationally, human THP-1 experiments provide proof of principle that this high-capacity, binucleated state can be induced in human macrophage model. Because the developmental functions of PDCD6IP, systemic perturbation is unlikely to be therapeutically feasible. More selective strategies could include engineering macrophages *ex vivo* to generate a high-capacity efferocytic state for adoptive transfer, targeting PDCD6IP specifically to lesional macrophages, or identify downstream transcriptional, metabolic, or secreted programs that reproduce the beneficial clearance and pro-resolving features of the binucleated state.

In summary, our work unexpectedly linked cytokinesis machinery to macrophage functional specialization, revealing an exciting biology where failed cell division acts as a pathway to generate a distinct, efferocytosis-specialized macrophage state that drives disease protection. Moving forward, harnessing this morphologically and transcriptomically unique macrophage state represents a compelling open opportunity to restore defective efferocytosis.

## DATA AVAILABILITY

All data are included in the manuscript or the Supplementary Information. The raw numbers for charts and graphs, and unedited gel images will be provided as the Source Data file. Additional raw data are available from the authors, as are unique reagents used in this article. The RNA-seq and scRNA-seq data will be deposited in the Gene Expression Omnibus (GEO) repository.

## CODE AVAILABILITY

Code for RNA-seq and scRNA-seq data analysis will be available via the Zhang lab GitHub repository (https://github.com/hanruizhang/).

## ACKNOWLEDGEMENTS

The authors’ research work has received funding from the National Institutes of Health (NIH) (R01HL151611, R01HL168174, R35HL177389), the M. Iréne Ferrer Scholar Award, and the Schaefer Research Scholar Award (to H.Z.), NIH P01HL172741 (to H.Z., I.T., A.R.T.), R35HL145228 and R35HL183005 (to I.T.), R01HL155431, R01HL107653, R01HL170157 (to A.R.T.), the American Heart Association Career Development Award 25CDA1451366 (to X.W.) and 23CDA1052177 (to F.L.).

We would like to acknowledge the NIH funding sources to the Columbia Stem Cell Initiative Flow Cytometry core facility by grant number S10OD036289, Columbia Center for Translational Immunology (CCTi) Flow Cytometry Core by grant number S10OD020056 and P30DK063608, the NIH/NCI P30CA013696 for the resources at the Herbert Irving Comprehensive Cancer Center Confocal and Specialized Microscopy Shared Resource, Molecular Pathology Shared Resource, and the Columbia Genome Center.

## AUTHOR CONTRIBUTIONS STATEMENT

X.W. and H.Z. conceived and designed the project. X.W. performed majority of the experiments. Z.W. performed *in vivo* efferocytosis assays. C.X. performed bioinformatic analyses. J.Y., T.S., J.C., and C.W. assisted experiments. F.L., W.L., and M.K. provided technical support. L.M., I.T. and A.R.T. provided critical intellectual input. X.W. and H.Z. drafted the manuscript. L.M., I.T. and A.R.T. critically read the manuscript. H.Z. supervised the project and funding. All the authors have read the manuscript and provided input to the manuscript.

## DECLARATION OF GENERATIVE AI AND AI-ASSISTED TECHNOLOGIES IN THE MANUSCRIPT PREPARATION PROCESS

During the preparation of this work, the author(s) used ChatGPT for checking grammar and improving clarity. The authors reviewed and edited the output as needed and take full responsibility for the content of the published article.

## METHODS

The Methods are organized to follow the experimental flow of the study: animal models; bone-marrow derived macrophage (BMDM) differentiation, apoptotic cell preparation, and *in vitro* efferocytosis assays by flow cytometry and microscopy-based analyses; single-cell RNA-sequencing (scRNA-seq) and data analyses; *in vivo* apoptotic cell clearance, autoimmunity model, and atherosclerosis studies; *ex vivo* splenic macrophage assays and whole-spleen bulk RNA-seq; and THP-1-based human cell validation experiments.

### Experimental animals

All animal procedures were performed in accordance with protocols approved by the Institutional Animal Care and Use Committee at Columbia University (Protocol #: AC-AABN5558). Mice were socially housed under specific pathogen-free conditions at 22 °C with 40–60% humidity on a 12 h light/12 h dark cycle with ad libitum access to food and water (PicoLab Rodent Diet 20 5053 and 5058, LabDiet). The Western diet (WD; TD88137, Envigo Teklad) was used for atherosclerosis study.

Heterozygous *Pdcd6ip* knockout mice (C57BL/6NCrl-*Pdcd6ipem1*(IMPC)Mbp/Mmucd; #043642) were obtained from the Knockout Mouse Project (KOMP) Repository through the Mutant Mouse Resource & Research Centers. Homozygous *Pdcd6ip* knockout mice and wild-type littermate controls were generated by intercrossing heterozygous *Pdcd6ip* knockout mice. Both male and female mice were used at 8-to 12-week-old unless otherwise specified, and experimental and control mice were co-housed.

### Bone marrow-derived macrophage differentiation

Bone marrow-derived macrophages (BMDMs) were generated as previously described^31^. Briefly, bone marrow cells were isolated from femurs and tibias by flushing with DMEM basal medium (Thermo Fisher Scientific, 11965092) using 10 mL syringes fitted with 26 G needles on day 0. Isolated bone marrow cells were cultured at 37 °C with 5% CO_2_ on non-tissue-culture-treated dishes in BMDM differentiation medium containing DMEM supplemented with 10% (vol/vol) heat-inactivated fetal bovine serum (HI-FBS; Thermo Fisher Scientific, A5209501), 20% (vol/vol) L-929 fibroblast-conditioned medium, and 2 mM L-glutamine (Thermo Fisher Scientific, 25030081). The first medium change was performed on day 4 or 5, and the medium was subsequently replaced every 2-3 days. BMDMs were considered fully differentiated and used for *in vitro* assays between days 7 and 9 after seeding.

### Preparation and fluorescent labeling of apoptotic cells

Apoptotic cells (ACs) were generated by exposing isolated mouse thymocytes to ultraviolet light (UV, 254 nm wavelength) irradiation using a UVP EL Series UV Lamp (Analytik Jena US, Upland, CA; Model UVS-18, 95-0200-01)^21^. Briefly, single-cell suspensions were prepared from thymi of C57BL/6J wild-type mice and seeded in 10 cm non-tissue-culture-treated dishes at 1 × 10^7^ cells per mL in 8 mL of 1× Dulbecco’s phosphate-buffered saline without calcium and magnesium (DPBS; Fisher Scientific, MT21031CM). Cells were irradiated for a total of 15 min and then incubated at 37 °C with 5% CO_2_ for 2.5 h. Fluorescent labeling of ACs was performed as described^20^. For single labeling, ACs were incubated with 2 μM Hoechst 33342 (Thermo Fisher Scientific, 62249) for 30 min or 10 μg/mL TAMRA (Thermo Fisher Scientific, C1171) for 25 min^20^. For dual labeling with Hoechst 33342 and pHrodo-Red (Thermo Fisher Scientific, P36600), ACs were first stained with 2 μM Hoechst 33342 for 30 min, followed by staining with 20 ng/mL pHrodo-Red for 25 min^31^. After staining, ACs were washed with a 10-fold volume of DPBS before use in efferocytosis assays.

### *In vitro* efferocytosis assay

As described^31^, bone marrow cells were seeded at 2 × 10^6^ cells per well of 6-well non-tissue culture-treated plates for differentiation into BMDMs. For a successful differentiation, we should expect to obtain 0.8–1.0 × 10^6^ BMDMs per well of 6-well plate, and the cells should appear ∼80% confluent. BMDMs are ready for efferocytosis assay on day 7. For flow cytometry-based analysis, fluorescently labeled ACs were co-incubated with macrophages at a 5:1 AC-to-macrophage ratio for 45 min in 2 mL of DMEM basal medium supplemented with 10% HI-FBS at 37 °C and 5% CO□. BMDMs were then gently washed five times with DPBS to remove unbound ACs and collected. For microscopy-based analysis, a lower AC-to-macrophage ratio of approximately 1:1 was used to reduce the number of free or unbound ACs in the imaging field, thereby minimizing background and facilitating the image acquisition and quantification of efferocytes.

### Flow cytometry-based quantification of efferocytosis and cargo acidification

For flow cytometry-based quantification of efferocytosis in BMDMs, cells were detached using CellStripper (Corning, 25-056-CI), a non-enzymatic cell dissociation solution, for live-cell analysis. Engulfment of ACs by macrophages was quantified as the percentage of macrophages positive for Hoechst-labeled AC cargo (Hoechst^+^ macrophages). Acidification of engulfed AC cargo was quantified as the percentage of pHrodo^+^ macrophages within the Hoechst^+^ macrophage population.

### Plasmid construction and lentiviral packaging for *PDCD6IP* overexpression

To determine the effects of PDCD6IP overexpression on macrophage efferocytosis, plasmid expressing human *PDCD6IP* cDNA was obtained from Addgene (#89859) and cloned into the pLE4-GFP lentiviral backbone^51^ to express PDCD6IP fused with GFP. PDCD6IP is highly conserved between humans and mice, including its major functional domains and protein-interaction regions. The use of the human construct also enabled exogenous PDCD6IP-GFP to be distinguished from endogenous murine expression. Lentiviral particles were generated in HEK293T cells (ATCC, CRL-3216) by co-transfection of the lentiviral expression vector with the packaging plasmid psPAX2 (Addgene, #12260) and the envelope plasmid pMD2.G (Addgene, #12259) using FuGENE 6 transfection reagent (Promega). The medium was replaced 16-18 h after transfection. Twenty-four hours after medium change, lentiviral supernatants were harvested and stored at 4 °C. Fresh medium was then added, and a second harvest of lentiviral supernatant was collected 24 h later. The two harvests were pooled and filtered through 0.45 μm SFCA filters (Corning, CLS431231).

### Lentiviral transduction of murine bone marrow cells for *PDCD6IP* overexpression

To assess the effects of *PDCD6IP* overexpression in BMDMs on efferocytosis, murine bone marrow cells were transduced with lentiviral vectors encoding either PDCD6IP-GFP or GFP alone as the control. On day 0, 8 × 10□ bone marrow cells were seeded in 10 cm Petri dish in a total volume of 10 mL, consisting of 3 mL of filtered crude lentiviral supernatant and 7 mL of BMDM differentiation medium supplemented with polybrene at a final concentration of 10 μg/mL (R&D Systems, 7711/10). Cells were maintained under standard BMDM differentiation conditions, and the culture medium was replaced on day 5. On day 7, BMDMs were replated to non-tissue-culture-treated 6-well plates. On day 8, efferocytosis assay was performed and assessed by flow cytometry. Because transgene expression was restricted to GFP-positive cells, all downstream analyses were performed after gating on the GFP-positive cell population.

### Macrophage binucleation and lysosomal imaging

For assessment of macrophage binucleation, macrophages were fixed with 4% formaldehyde (Thermo Fisher Scientific, 28908) for 30 min, permeabilized with 0.1% Triton X-100 in DPBS for 10 min, and stained with 2 μM Hoechst 33342 and Alexa Fluo 488 conjugated phalloidin (Phalloidin-AF488; Invitrogen, A12379, 1:500) for 15 min. For centrosome staining, blocking was performed using 5% BSA in DPBS. Macrophages were incubated with anti-pericentrin antibody (Abcam, ab4448, 1:50) overnight at 4°C, followed by goat anti-rabbit Alexa Fluor 647 secondary antibody (Invitrogen, A-21245, 1:200). Mononuclear and binucleated macrophages were identified by Nikon spinning-disk confocal microscope equipped with a 60x/1.49 Apo TIRF oil objective.

For lysosome imaging, live macrophages were stained with LysoTracker (Invitrogen, L7528, 1:1000) for 20 min at 37 °C with 5% CO_2_. Cells were then washed with DPBS containing 2% HI-FBS and nuclei stained with DRAQ5 (Invitrogen, 65-0880-92, 1:2000) before imaging by Nikon spinning-disk confocal microscope equipped with a 60x/1.49 Apo TIRF oil objective. LysoTracker fluorescence intensity per cell was quantified using ImageJ v1.54p (NIH).

### Microscopy-based quantification of AC binding, engulfment, and time required for the completion of engulfment in bone-marrow derived macrophages

The effects of *Pdcd6ip* knockout on AC binding, engulfment, and time required for the completion of engulfment were assessed in BMDMs. To quantify AC binding, macrophages were pretreated with 5 µM cytochalasin D (Sigma-Aldrich, Cat# C8273), an actin polymerization inhibitor, for 45 min to block internalization. Macrophages were then incubated with TAMRA-labeled ACs for 15 min. Unbound ACs were removed by washing with DPBS, fixed, stained for nuclei (DRAQ5) and F-actin (Phalloidin-AF488), and imaged. To quantify AC engulfment, macrophages were incubated with TAMRA-labeled ACs for 45 min, followed by washing to remove unbound ACs. Cells were then fixed, stained for nuclei (DRAQ5) and F-actin (Phalloidin-AF488) and imaged. Imaging was performed using Nikon spinning-disk confocal microscope equipped with a 60x/1.49 Apo TIRF oil objective.

To determine the time required for completion of efferocytosis, time-lapse live-cell imaging was performed using a Nikon spinning-disk confocal microscope equipped with a 60x/1.49 Apo TIRF oil objective. Macrophages were stained with LysoTracker (Invitrogen, L7528, 1:1000) for 20 min at 37 °C with 5% CO_2_, to facilitate visualization of macrophage morphology during live cell imaging. Cells were then washed with DPBS containing 2% HI-FBS and nuclei stained with DRAQ5 (Invitrogen, 65-0880-92, 1:2000). ACs were labeled by Hoechst 33342. ACs were added to macrophages at a 1:1 AC-to-macrophage ratio. Images were then acquired every 3 min for a total of 15 min.

### Western blotting of cell lysate

Cells from one well of a six-well plate, corresponding to approximately 1 × 10^6^ cells, were harvested using CellStripper (Corning, 25-056-CI) and pelleted by centrifugation. Cell pellets were lysed on ice for 30 min in 70 μL of 1× RIPA buffer (Sigma-Aldrich, 20-188) supplemented with protease inhibitor cocktail (Roche, 04574834001), followed by centrifugation at 12,000 × g for 10 min at 4 °C, and the supernatants were transferred to fresh tubes. Lysates were mixed with 5× SDS sample buffer containing 250 mM Tris-HCl (pH 6.8), 20% SDS, 30% glycerol (v/v), 10 mM DL-dithiothreitol (DTT), and 0.05% bromophenol blue (w/v). Samples were loaded onto 4–20% Tris-Glycine gels (Invitrogen, EC6021BOX) and transferred onto 0.2 μm PVDF membranes (Thermo Fisher Scientific, 88520). Membranes were blocked with 5% nonfat dry milk in TBST (Thermo Fisher Scientific, J77500.K8) for 1 h at room temperature. Primary antibody incubation was performed at 4 °C overnight. After washing with TBST, membranes were incubated with appropriate secondary antibodies for 1 h at room temperature. The primary antibody used was anti-PDCD6IP (Cell Signaling Technology, 2171S, 1:1000) followed by incubation with HRP-linked secondary antibody (Cell Signaling Technology, 7076). HRP-conjugated rabbit anti-β-actin (Cell Signaling Technology, 5125S, 1:2000) was used as a loading control. After the final wash to remove unbound antibodies, the protein expression was detected by SuperSignalTM West Pico PLUS Chemiluminescent Substrate (Thermo Fisher Scientific, 34580) and imaged using ChemiDoc Imaging System (Bio-Rad). Band intensity was quantified using ImageJ v1.53u (NIH).

### Western blotting of conditioned medium

For detecting secreted protein, 100 μL of conditioned medium was mixed with 5× SDS sample buffer. Samples were separated on 7% Tris-acetate gels (Invitrogen, EA0358BOX) and transferred onto 0.2 µm PVDF membranes (Thermo Fisher Scientific, 88520) using a Bio-Rad semi-dry transfer system (Bio-Rad, 1703940) for 15 min. Membranes were stained with Ponceau S staining solution (Thermo Fisher Scientific, A40000279) for 3-5 min and scanned using an HP LaserJet Pro MFP M426fdn Printer (HP, F6W14A#BGJ). Ponceau S staining solution was removed by washing the membranes once with TBST for 5 min. Membranes were then blocked with 5% nonfat dry milk in TBST for 30 min, washed once with TBST for 5 min, and incubated overnight at 4 °C with an anti-TGF-β1 antibody (HUABIO, HA721143; 1:500) followed by incubation with HRP-link secondary antibody (Cell Signaling Technology, 7074).

### BD Discover S8 imaging cytometry

Image-enabled flow cytometric analysis was performed using a BD FACSDiscover S8 Cell Sorter equipped with BD CellView Image Technology (BD Biosciences). BMDMs were stained with Vybrant DyeCycle Green (Invitrogen, V35004; 1:5000) to assess DNA content. Because DNA content alone cannot distinguish binucleated cells from mononuclear cells with replicated DNA content, image-based nuclear morphology was used to classify cells as mononuclear or binucleated. Data were acquired using BD FACSChorus software v6.2.0 and analyzed using FlowJo v10 with BD CellView Lens plugin 1.2.3. Automated image-based scoring was followed by manual inspection of representative cell images to confirm classification accuracy. Cells with weak DyeCycle Green signal, poor image quality, or ambiguous nuclear morphology that prevented confident assignments as mononuclear or binucleated were excluded from scoring. This approach was used as an independent imaging flow cytometry-based method to validate macrophage binucleation and its relationship with DyeCycle Green-based DNA content measurement, complementing microscopy-based quantification.

### Sorting of mononuclear and binucleated BMDMs for single-cell RNA-sequencing

To maximize post-sorting viability, BMDMs were sorted using a BD FACSAria cell sorter (BD Biosciences) forscRNA-seq analyses. BMDMs were stained with Vybrant DyeCycle Green (Invitrogen, V35004; 1:1000). Based on the gating strategy established by imaging flow cytometry, DyeCycle Green^hi^ FSC-A^hi^ cells were sorted to enrich binucleated macrophages. DyeCycle Green^low^ FSC-A^low^ cells were sorted to enrich monunuclear macrophages. Mononuclear and binucleated cell-enriched populations were sorted separately, with a target of 10,000 cells collected from each population. Enrichment of mononuclear or binucleated cells in the sorted populations was confirmed by reserving an aliquot from each population for fluorescence microscopy. The remaining cells were subjected to cell hashing using TotalSeq-B hashtags. Mononuclear cell-enriched *Pdcd6ip^+/+^* BMDMs, *Pdcd6ip^+/+^* (mono), were labeled with Hashtag #1/TotalSeq-B0301, barcode ACCCACCAGTAAGAC (BioLegend, 155831); Mononuclear cell-enriched *Pdcd6ip^−/−^* BMDMs, *Pdcd6ip^−/−^* (mono), were labeled with Hashtag #2/TotalSeq-B0302, barcode GGTCGAGAGCATTCA (BioLegend, 155833); and binucleated cell-enriched *Pdcd6ip^−/−^* BMDMs, *Pdcd6ip^−/−^* (bi), were labeled with Hashtag #3/TotalSeq-B0303, barcode CTTGCCGCATGTCAT (BioLegend, 155835). Live cells were submitted to the Single Cell Analysis Core at the J. P. Sulzberger Columbia Genome Center for two 10x Chromium scRNA-seq runs.

### scRNA-seq data processing and analysis

Reads from *Pdcd6ip^+/+^* (mono), *Pdcd6ip^−/−^*(mono), and *Pdcd6ip^−/−^* (bi) populations were processed and demultiplexed using Cell Ranger 9.0.0 multi pipeline with GENCODE vM23 annotation from pre-built mouse reference package “mm10-2020-A”. Gene and cell level filtering were performed in each demultiplexed sample. Genes that were expressed in <10 cells were excluded from analysis. Cells were retained if 1) ≥200 genes were expressed, and 2) percentage of reads mapped to mitochondrial genes was <10%. In addition, an upper limit of 5,500 genes and 40,000 UMIs were applied to cells from *Pdcd6ip^+/+^* (mono) and *Pdcd6ip^−/−^*(mono) samples. Due to higher RNA content in binucleated cells, an upper limit of 6,500 genes and 65,000 UMIs were applied to the *Pdcd6ip^−/−^*(bi) sample. Clustering analysis was performed in CarDEC^52^, a deep learning framework for batch effect correction, gene expression denoising and clustering. A total of 2,000 highly variable genes was selected. No batch variable was set, given minimal batch effect between multiplexed samples in the same sequencing run. A total of 6 clusters were identified that reflected known macrophage population and did not show over-clustering. Differential expression analysis was conducted in Scanpy 1.8.1^53^. Raw counts were normalized by the total counts per cell using a target sum of 10,000 and then log-transformed. Gene expression from each cluster was compared to cells in other clusters using Wilcoxon rank-sum test. P-values were adjusted by Benjamini-Hochberg approach. Differentially expressed (DE) genes were defined by an absolute fold change ≥1.5, adjusted P-value <0.05, and expression in ≥25% of cells from either group. Cell types were annotated by comparing the top DE genes to literature.

Cell composition analysis: To compare cell type abundance changes in *Pdcd6ip^−/−^* (mono) and *Pdcd6ip^−/−^* (bi) populations compared to the *Pdcd6ip^+/+^* (mono) population, we applied scCODA^54^, a Bayesian model for cell type compositional analysis in single cell data that controls for false discovery rate (FDR) with small sample size. Counts from each group and each cell type were tabulated as input to scCODA. The “group” variable was included as the covariate and the reference cell type was set to “automatic”, which selected “Cluster 3” as the reference as it had minimal change in proportion between groups. FDR level was set to 0.2 as the default level of 0.05 was not able to detect credible effects given small sample size.

### Pseudobulk differential expression and pathway analysis

To compare gene expression between individual populations, we applied pseudobulk approach on scRNA-seq data for robust differential expression analysis. Cells from each sample were randomly sampled to generate 5 technical replicates. Read counts were summed across cells from the sample technical replicate to generate pseudobulk counts. The pseudobulk approach was applied to all cells as one sample. Differential expression analysis was conducted using DESeq2 1.42.0^55^. Genes were filtered if the total pseudobulk counts across replicates was <50. Log2 fold changes were shrunken using apeglm method^56^ implemented in DESeq2. P-values were adjusted by Benjamini-Hochberg approach. Genes with an absolute fold change ≥1.5 and adjusted P-value <0.05 were considered DE genes. Over-representation analysis (ORA) was performed in clusterProfiler 4.10.0^57^ to identify enriched gene sets from DE genes using Gene Ontology (Biological Process, Cellular Component, and Molecular Function). In all ORA analyses, the background gene set was chosen as all genes that passed pseudobulk count filtering in the corresponding DESeq2 analysis. P-values were adjusted using Benjamini-Hochberg approach. Gene sets with adjusted P-value <0.05 were considered significantly over-represented. Enriched terms from Gene Ontology analysis were simplified using “simplify” function to remove redundancy.

### Assessment of TAMRA-labeled apoptotic cell uptake by splenic macrophages *in vivo*

ACs were labeled with 10 μg/mL TAMRA (Thermo Fisher Scientific, C1171) for 25 min. A total of 2 × 10^7^ ACs per mouse were administered intravenously via tail vein injection^21^. At 24 or 48 h post-injection, mice were euthanized, and the spleen was harvested. Single-cell suspensions were prepared from the spleen using the Spleen Dissociation Kit (Miltenyi Biotec, 130-095-926), according to the manufacturer’s instructions. The percentage of TAMRA^+^ splenic macrophages was determined by flow cytometry. Splenic macrophages were identified using the following antibody cocktail: CD45.2-PerCP/Cyanine5.5 (BioLegend, 109828, 1:50), CD11b-APC (BioLegend, 101211, 1:50), and F4/80-FITC (BioLegend, 123108, 1:50).

### Assessment of autoimmunity-related phenotypes in mice after repeated apoptotic cell injections

Exposure to large numbers of ACs was used to induce autoimmune responses in mice, as previously described^21^. Thymocytes from C57BL/6J wild-type mice were UV-irradiated to induce apoptosis, and 2 × 10^7^ ACs per mouse were administered intravenously via tail vein injection. Injections were performed weekly for 4 weeks, followed by a 2-week pause and two additional injections. One week after the final injection, mice were euthanized, and plasma and kidneys were collected to assess autoimmune responses.

Plasma autoantibodies: Plasma antinuclear antibodies (ANA) and anti-dsDNA antibodies were measured using a mouse antinuclear antigens Ig’s total IgA/IgG/IgM ELISA kit (Alpha Diagnostics International, 5210), and a mouse anti-dsDNA Ig’s total IgA/IgG/IgM ELISA kit (Alpha Diagnostics International, 5110).

Immune complex deposition in glomeruli: Kidneys were embedded in optimal cutting temperature (OCT) compound and snap frozen. Cryostat sections were cut at 3 μm. For immunofluorescence staining of C1q, IgG, or wheat germ agglutinin (WGA), frozen sections were fixed with 4% formaldehyde for 20 min at room temperature. Sections were permeabilized with 0.5% Triton X-100 in PBS and blocked with 5% BSA in PBS containing 0.05% Tween-20 (PBST). Slides were washed three to five times with PBST between steps. For C1q staining, sections were incubated overnight at 4 °C with rabbit monoclonal anti-C1q antibody (Abcam, ab182451, 1:50), followed by incubation for 1 h at room temperature with Alexa Fluor 555-conjugated goat anti-rabbit IgG (H+L) secondary antibody (Invitrogen, A-21428, 1:200). For IgG and WGA staining, Alexa Fluor 555-conjugated anti-IgG antibody (Invitrogen, A-21422, 1:200) and Alexa Fluor 488-conjugated WGA (Invitrogen, W11261, 1:5000) were used, respectively. DAPI (Invitrogen, P36931, 1:5000) was used to stain nuclei. Negative controls were included using rabbit IgG isotype control antibody (Cell Signaling Technology, 3900S, 1:50). Slides were mounted with antifade mountant (Invitrogen, P36930) and imaged using the ImageXpress Micro4 high-content microscopy with a Nikon Plan Apo λ 20×/0.75 objective (Molecular Device) or Nikon Ti-S Automated Inverted Microscope (Nikon). The mean fluorescence intensity (MFI) within the glomeruli area was quantified using ImageJ v1.53u (NIH).

### Bone marrow transplantation and atherosclerosis studies

To achieve hematopoietic *Pdcd6ip* knockout, *Ldlr^−/−^*mice (#002207, the Jackson Laboratory) were lethally irradiated and then transplanted with bone marrow from homozygous *Pdcd6ip* knockout (*Pdcd6ip^−/−^*) mice or wild-type (*Pdcd6ip^+/+^*) littermate controls. Bone marrow cells were isolated from the femurs and tibias of *Pdcd6ip^+/+^*and *Pdcd6ip^−/−^* donor mice. Male and female *Ldlr^−/−^*recipient mice aged 12-13 weeks were lethally irradiated with a single dose of 10.5 Gy^58^ using a MultiRad 350 X-ray irradiator (Precision X-Ray, USA) operated at 350 kVp and 11.4 mA with an Sn/Cu/Al filter at a dose rate of 1.22 Gy/min. Within 5 h after irradiation, bone marrow was collected from donor mice aged 10-12 weeks with the indicated genotypes. Irradiated recipient mice were randomized to experimental groups and received 6 × 10^6^ total bone marrow cells by tail vein injection. After 5 weeks of reconstitution, all bone marrow-transplanted mice were fed a Western diet (WD; TD88137, Envigo Teklad) for 21-23 weeks. Total plasma cholesterol was measured using a FUJIFILM Free Cholesterol E kit (Fisher Scientific, NC9506034).

### Complete blood cell count (CBC) and differential count

Retro-orbital bleeding was performed to collect ∼200□μL blood per mouse using BD Microtainer tubes with K2EDTA (BD, 365974) for complete blood count and differential count using a Heska Element HT5 by the diagnostic lab at the Institute of Comparative Medicine, Columbia University Irvine Medical Center^59^.

### Histological quantification of aortic root lesions

At the study end point, mice were euthanized and hearts were collected and fixed in 10% formalin for 24 h. Fixed heart tissues were embedded in paraffin and serially sectioned. Serial sections of the aortic root were collected from the appearance of the three aortic valve leaflets through the disappearance of the three valve leaflets toward the ascending aorta. Sections were cut at 5 µm thickness and six sections spaced 30 µm apart toward the ascending aorta were selected for hematoxylin and eosin (H&E) staining. The average lesion area and necrotic core area from the selected sections were calculated for each mouse and used to determine lesion size and necrotic core area. Necrotic core was defined as an acellular area devoid of intact cells^60^. All sections were scanned at ×40 magnification using a Leica AT2 whole-slide digital imaging system. Imaging analysis was performed using Aperio ImageScope software (Leica, v12.4.6.5003).

### Fibrous cap thickness analysis

Aortic root sections (one section per mouse) obtained from a comparable anatomical location across mice with relatively larger lesion size were stained with Picrosirius Red according to the manufacturer’s instructions (Polysciences, 24901-500). All sections were scanned at ×40 magnification using a Leica AT2 whole-slide digital imaging system. Imaging analysis was performed using Aperio ImageScope software (Leica, v12.4.6.5003). The average thickness from all measurements within each section was calculated and reported for each mouse.

#### *In situ* plaque efferocytosis assay^61^

Paraffin-embedded tissue sections were incubated at 55 °C and deparaffinized at the Columbia Molecular Pathology Shared Resource core. Following deparaffinization and rehydration, heat-induced antigen retrieval was performed using Trilogy Pretreatment Solution (Cell Marque, 920P-09). Sections were permeabilized with 0.5% Triton X-100 in PBS, and subjected to TUNEL staining according to the manufacturer’s instructions (Invitrogen, C10619). After TUNEL labeling, sections were blocked with 5% BSA in PBST for 30 min and incubated overnight at 4°C with an anti-CD68 antibody (Abcam, ab53444, 1:100), followed by a goat anti-rat Alexa Fluor 647 secondary antibody (Invitrogen, A48265TR; 1:200). Nuclei were counterstained with DAPI. Images were acquired using a Nikon Ti-S Automated Inverted Microscope. For quantification, TUNEL^+^ nuclei in close proximity to or in contact with CD68^+^ macrophages were counted as macrophage-associated ACs, indicative of efferocytosis. TUNEL^+^ nuclei without neighboring macrophages were counted as free ACs. The ratio of macrophage-associated ACs to free ACs was calculated as a measure of lesional efferocytosis^61^.

#### Immunofluorescence staining of aortic root lesions

Paraffin sections were incubated at 55 °C and deparaffinized at the Columbia Molecular Pathology Shared Resource core. For antigen retrieval, paraffin-embedded slides were rehydrated in Trilogy solution (Cell Marque, 920P-09). The following primary antibodies were used for staining: anti-TGF-β1 (HUABIO, HA721143; 1:100) and anti-IL-1β (HUABIO, HA601036; 1:100). Primary antibodies were incubated with the sections overnight at 4°C. The following secondary antibodies were used: goat anti-rabbit Alexa Fluor 555 secondary antibody (Invitrogen, A-21428; 1:200), and goat anti-mouse Alexa Fluor 555 secondary antibody (Invitrogen, A-21424; 1:200). Sections were imaged using a Nikon Ti-S Automated Inverted Microscope. Fluorescence intensity in plaque was quantified using ImageJ v1.54p (NIH).

#### *Ex vivo* splenic macrophage isolation and efferocytosis

*Ldlr*^−/−^ recipient mice were lethally irradiated and transplanted with bone marrow from *Pdcd6ip^+/+^* or *Pdcd6ip^−/−^*donor mice. After 5 weeks of reconstitution, mice were fed a Western diet for 21-23 weeks, as described for atherosclerosis studies. At the study endpoint, mice were euthanized and spleens were harvested. Spleens were dissociated using the Spleen Dissociation Kit (Miltenyi Biotec, 130-095-926), according to the manufacturer’s instructions. The dissociated tissue was resuspended in 5 mL of DPBS supplemented with 2% HI-FBS, 5□mM EDTA (Thermo Fisher Scientific,15-575-020), 1□mM sodium pyruvate (Fisher Scientific, 11-360-070), and 20□mM HEPES (Fisher Scientific, MT25060CI). For positive selection of CD11b^+^ splenic macrophages, 100 μL of CD11b MicroBeads (Miltenyi Biotec, 130-097-142) was added to each 5 mL cell suspension, followed by incubation for 30 min at 4 °C. CD11b^+^ splenic macrophages were isolated by positive selection using magnetic separation according to the manufacturer’s instructions. Purified cells were seeded at 5,000-10,000 cells per confocal dish (Alkali Scientific TCG22) in BMDM differentiation medium and incubated for 6-8 h to allow adherence before analyses.

To quantify the percentage of binucleated splenic macrophages, macrophages were stained with DRAQ5 (Invitrogen, 65-0880-92, 1:2000) and imaged live cell using a Nikon spinning-disk confocal microscope equipped with a 60x/1.49 Apo TIRF oil objective. To quantify AC engulfment, macrophages were incubated with TAMRA-labeled ACs at a 1:1 AC-to macrophage ratio for 45 min, followed by washing to remove unbound ACs. Cells were then fixed, stained for nuclei (DRAQ5) and F-actin (Phalloidin-AF488) and imaged. Imaging was performed using a Nikon spinning-disk confocal microscope equipped with a 60x/1.49 Apo TIRF oil objective.

### Bulk RNA-seq library preparation from spleen

*Ldlr*^−/−^ recipient mice were lethally irradiated and transplanted with bone marrow from *Pdcd6ip^+/+^* or *Pdcd6ip^−/−^*donor mice. After 5 weeks of reconstitution, mice were fed a Western diet for 21-23 weeks, as described for atherosclerosis studies. At the study endpoint, mice were euthanized and whole-spleen was harvested for bulk RNA-seq analysis. Total RNA extraction was performed using Monarch Spin RNA Isolation Kit (NEB, T2110S). RNA concentration and purity were assessed by NanoDrop 2000, and RNA integrity was evaluated using an Agilent TapeStation 4200; samples with RIN values greater than 7.0 were used for library preparation. RNA-seq was performed by IDseq/Innomics. Libraries were prepared using 200 ng total RNA. Briefly, polyadenylated mRNA was enriched using oligo(dT) magnetic beads, fragmented, reverse-transcribed into cDNA, end-repaired, A-tailed, ligated with MGI-specific adapters and sample barcodes, PCR-amplified, and purified. Indexed libraries were pooled at equimolar ratios, circularized, converted into DNA nanoballs, and sequenced on the DNBSEQ-G400 platform using paired-end 150 bp sequencing. Raw reads were demultiplexed, quality-checked using FastQC, and filtered to remove adapters, low-quality reads, and reads containing high levels of undefined bases before downstream analysis.

### Bulk RNA-seq data processing and differential expression analysis

Gene expression was quantified in each sample using a decoy-aware transcriptome and selective alignment approach in Salmon 1.10^62^. The decoy-aware transcriptome was built by concatenating GENCODE vM23 transcriptome and mouse genome build GRCm38, followed by Salmon indexing, specifying the chromosome names as the decoys. Reads were quantified using Salmon quant with GC bias correction. Transcript counts were imported using tximport package^63^ and aggregated to gene counts. Genes with total counts ≥10 across all samples were retained for differential expression analysis between *Pdcd6ip^−/−^*and *Pdcd6ip^+/+^* groups. Differential expression analysis followed the same process described in the pseudobulk analysis. DE genes were defined by an absolute fold change ≥1.5 and adjusted P-value <0.05.

### Plasmid construction and lentiviral packaging for CRISPR/Cas9-mediated *PDCD6IP* perturbation

Single-guide RNAs (sgRNAs) targeting human *PDCD6IP* gene were selected from the Human CRISPR Knockout Pooled Library (GeCKO v2) (Addgene, #1000000048). Candidate sgRNAs were evaluated and the most efficient sgRNA targeting *PDCD6IP* (5′-AGATGCCATCATAGCTAAAT-3′) was selected for subsequent experiments. The *PDCD6IP*-targeting sgRNA and a non-targeting control sgRNA (5′-ACACTCTTGTTCTTGCGCA-3′) were cloned into the pLenti H5070-Cas9 backbone. Plasmid construction, lentiviral packaging, and lentiviral particle concentration by ultracentrifugation were performed by OBiO Technology.

### THP-1 cell culture and lentiviral transduction of THP-1 cells for CRISPR/Cas9-mediated *PDCD6IP* perturbation and characterization of binucleation

Human THP-1 monocyte cell line (ATCC, TIB-202) was maintained in RPMI basal medium (Thermo Fisher Scientific, A4192301) supplemented with 10% HI-FBS, 1 mM sodium pyruvate, 10 mM HEPES, 50 μM 2-mercaptoethanol (Gibco, 21985023), and 2 mM L-glutamine (Coring, 25-005-CI)^20^. Cells were confirmed to be mycoplasma-free using the MycoAlert Mycoplasma Detection Kit (Lonza, LT07-418).

For CRISPR/Cas9-mediated gene perturbation, THP-1 monocytes were passaged and seeded at 1 × 10^5^ cells per well in 6-well plates. Immediately after seeding, cells were transduced with 10 μL concentrated lentivirus (OBiO Technology) encoding Cas9 and either a non-targeting control sgRNA or a *PDCD6IP*-targeting sgRNA in the presence of 10 μg/mL polybrene (R&D Systems, 7711/10). After transduction, cells were allowed to expand for 5 days before counted, passaged, and transduced again using the same procedure. This passaging and transduction workflow was repeated for three consecutive rounds. Cells were analyzed after the third round of transduction and expansion. Binucleation of transduced THP-1 monocytes was characterized by Nikon Ti-S Automated Inverted Microscope equipped with a Plan Apo λ 60×/1.40 oil objective, following nuclear staining using 2 μM Hoechst 33342 in live cells.

### THP-1 monocyte differentiation into macrophage-like cells, characterization of binucleation, *in vitro* efferocytosis, and TGF-β1 expression

THP-1 monocytes were seeded in 6-well plates at a density of 1 × 10^6^ cells per well and differentiated into macrophages by incubation with 100 nM phorbol 12-myristate 13-acetate (PMA; Sigma-Aldrich, P1585) in THP-1 culture medium for 24 h. The PMA-containing medium was then removed and replaced with fresh THP-1 culture medium, and the cells were cultured for an additional 48 h^20^.

To assess binucleation, differentiated THP-1 macrophages were stained using 2 μM Hoechst 33342 and subjected to live cell imaging. To assess efferocytosis, differentiated THP-1 macrophages were co-incubated with ACs for 45 min. Unbound ACs were removed by washing with DPBS. Cells were fixed, stained with 2 μM Hoechst 33342 and Phalloidin-AF488 to visualize nuclei and F-actin, respectively. To assess TGF-β1 expression, THP-1-derived macrophages were fixed with 4% formaldehyde for 30 min at room temperature, washed with DPBS, and permeabilized with 0.1% Triton X-100 in DPBS for 10 min. Cells were then blocked with 5% BSA in DPBS for 1 h at room temperature, followed by incubation with an anti-TGF-β1 primary antibody (HUABIO, HA721143; 1:100). After three washes with PBST, cells were incubated with goat anti-rabbit Alexa Fluor 555 secondary antibody (Invitrogen, A-21428; 1:200) for 1 h at room temperature. Cells were then stained for nuclei (Hoechst 33342) and F-actin (Phalloidin-AF488) and imaged. All images were acquired using a Nikon Ti-S Automated Inverted Microscope equipped with a Plan Apo λ 60×/1.40 oil objective.

### Statistical analysis

Statistical analyses were performed using GraphPad Prism 8. A two-tailed Student’s t-test was used to compare two groups. For one independent variable with more than two groups, one-way ANOVA was performed, followed by Tukey’s post-hoc test. For scenarios involving two independent variables (factors) with two or more groups, a two-way ANOVA was performed, followed by Bonferroni’s post-hoc test to adjust for multiple comparisons, as specified in the figure legends. Data are presented as mean□±□standard error of the mean (SEM). A P-value of 0.05 or less was considered statistically significant. The exact P values, as well as the number of independent experiments or biological replicates, are specified in figures and figure legends.

