## Supplementary for "Cytokinesis Arrest-induced Binucleation of Macrophages Produces Highly Efficient Efferocytes"

**SUPPLEMENTARY FIGUREs**

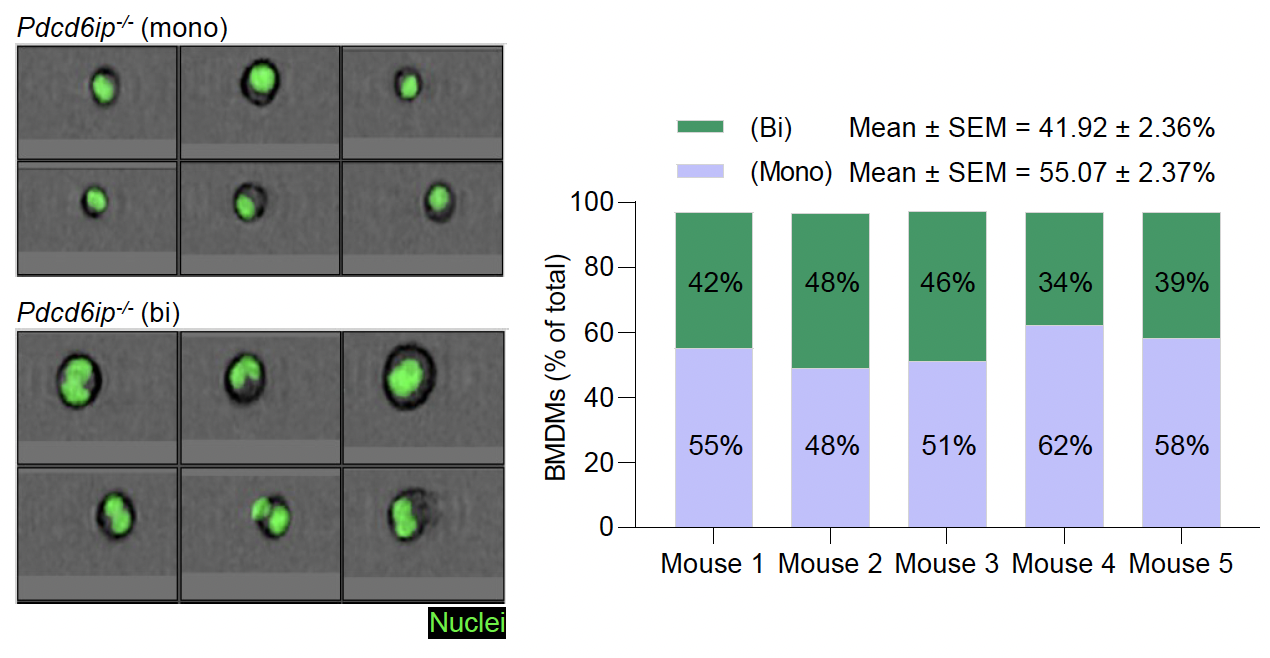

**Supplementary Fig. 1 | Imaging cytometry confirms increased binucleation in *Pdcd6ip^−/−^* macrophages.**

Bone marrow derived macrophages (BMDMs) from *Pdcd6ip^−/−^* mice were stained with DyeCycle Green to label DNA content and analyzed using a BD FACSDiscover S8 Cell Sorter. Nuclear numbers were determined from imaging cytometry images. Representative imaging cytometry images show mononuclear (mono) and binucleated (bi) BMDMs of *Pdcd6ip^−/−^* mice. Quantification shows the percentage of *Pdcd6ip^−/−^* (bi) BMDMs, consistent with microscopy-based analysis shown in **Fig. 2a**. Only cells with clearly resolved nuclear staining were scored, whereas cells with poor staining or ambiguous nuclear morphology were excluded from analysis; these cells account for ~3% of total images assessed. 200-300 evaluable cells were quantified per mouse. Data are shown as mean ± standard error of the mean (SEM). *n* = 5 *Pdcd6ip^−/−^* mice.

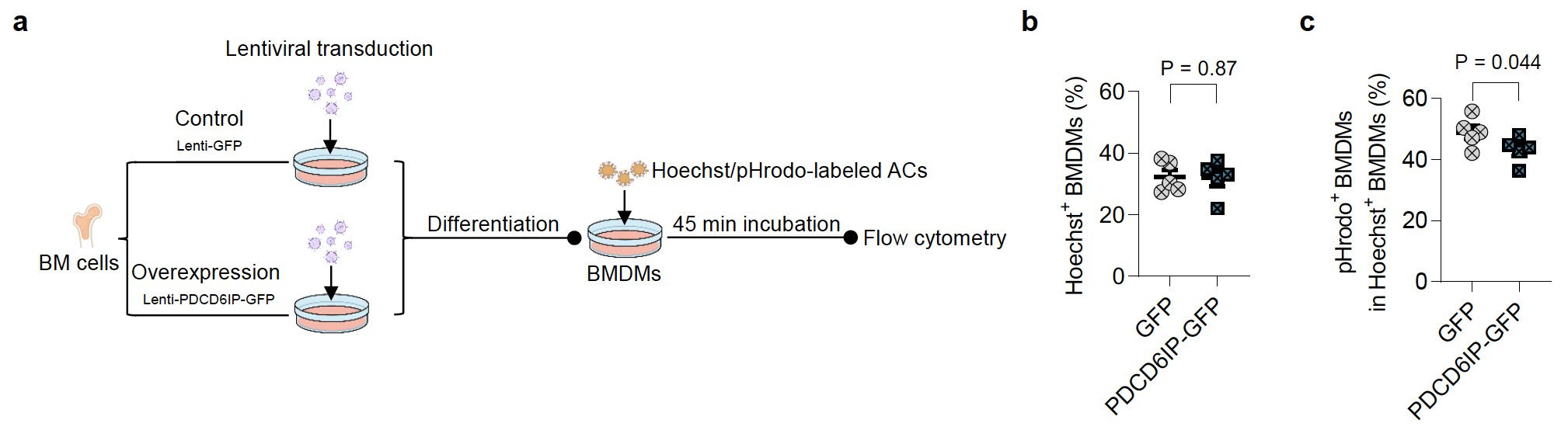

**Supplementary Fig. 2 | PDCD6IP overexpression does not alter macrophage engulfment but modestly reduces acidification in *in vitro* efferocytosis assay.**

**a,** Schematic of the experimental design. Bone marrow (BM) cells were transduced with lentiviral vectors expression GFP control or PDCD6IP-GFP, followed by differentiation into BMDMs. BMDMs were then used in an *in vitro* efferocytosis assay to assess apoptotic cell (AC) engulfment and cargo acidification. ACs were dual-labeled with Hoechst (pH-insensitive uptake marker) and pHrodo probe (pH-sensitive acidification maker), and co-incubated with BMDMs for 45 min. Hoechst^+^ BMDMs were quantified as AC-engulfing cells, whereas pHrodo positivity among Hoechst^+^ BMDMs was used to assess acidification of engulfed ACs. **b,** Quantification of Hoechst^+^ BMDMs. **c,** Quantification of pHrodo^+^ BMDMs among Hoechst^+^ BMDMs. *n* = 5 mice. Data are shown as mean ± SEM. Statistical significance was determined using a two-tailed paired Student’s *t*-test.

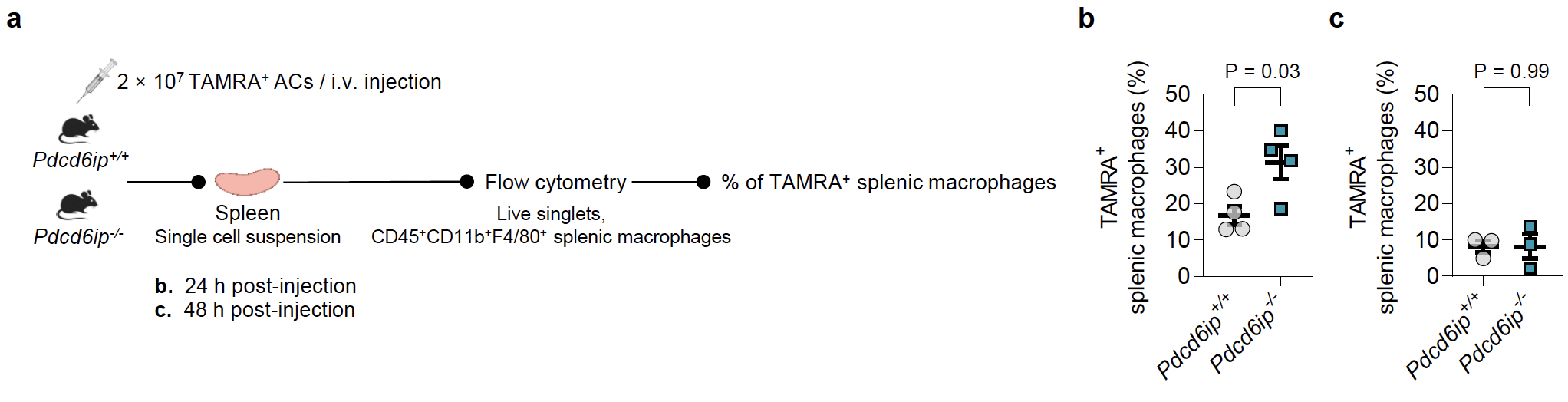

**Supplementary Fig. 3 | *Pdcd6ip* deletion enhances *in vivo* apoptotic cell clearance.**

**a,** Schematic of the experimental design. TAMRA-labeled ACs were injected into *Pdcd6ip*^+/+^ and *Pdcd6ip*^−/−^ mice intravenously via tail vein injection, and efferocytosis by splenic macrophages was quantified 24 and 48 h later by measuring TAMRA^+^ splenic macrophages (CD45^+^CD11b^+^F4/80^+^) using flow cytometry. **b,** Quantification of TAMRA^+^ splenic macrophages 24 h after AC injection, showing increased AC engulfment in *Pdcd6ip*^−/−^ mice. *n* = 4 *Pdcd6ip*^+/+^ and 4 *Pdcd6ip*^−/−^ mice. **c,** Quantification of TAMRA^+^ splenic macrophages 48 h after AC injection, showing reduced TAMRA positivity compared with 24 h after AC injection and no significant difference between *Pdcd6ip*^+/+^ and *Pdcd6ip*^−/−^ mice, consistent with degradation of engulfed cargos and clearance of the fluorescent signal. *n* = 3 *Pdcd6ip*^+/+^ and 3 *Pdcd6ip*^−/−^ mice. Data are shown as mean ± SEM. Statistical significance was determined using a two-tailed Student’s *t*-test.

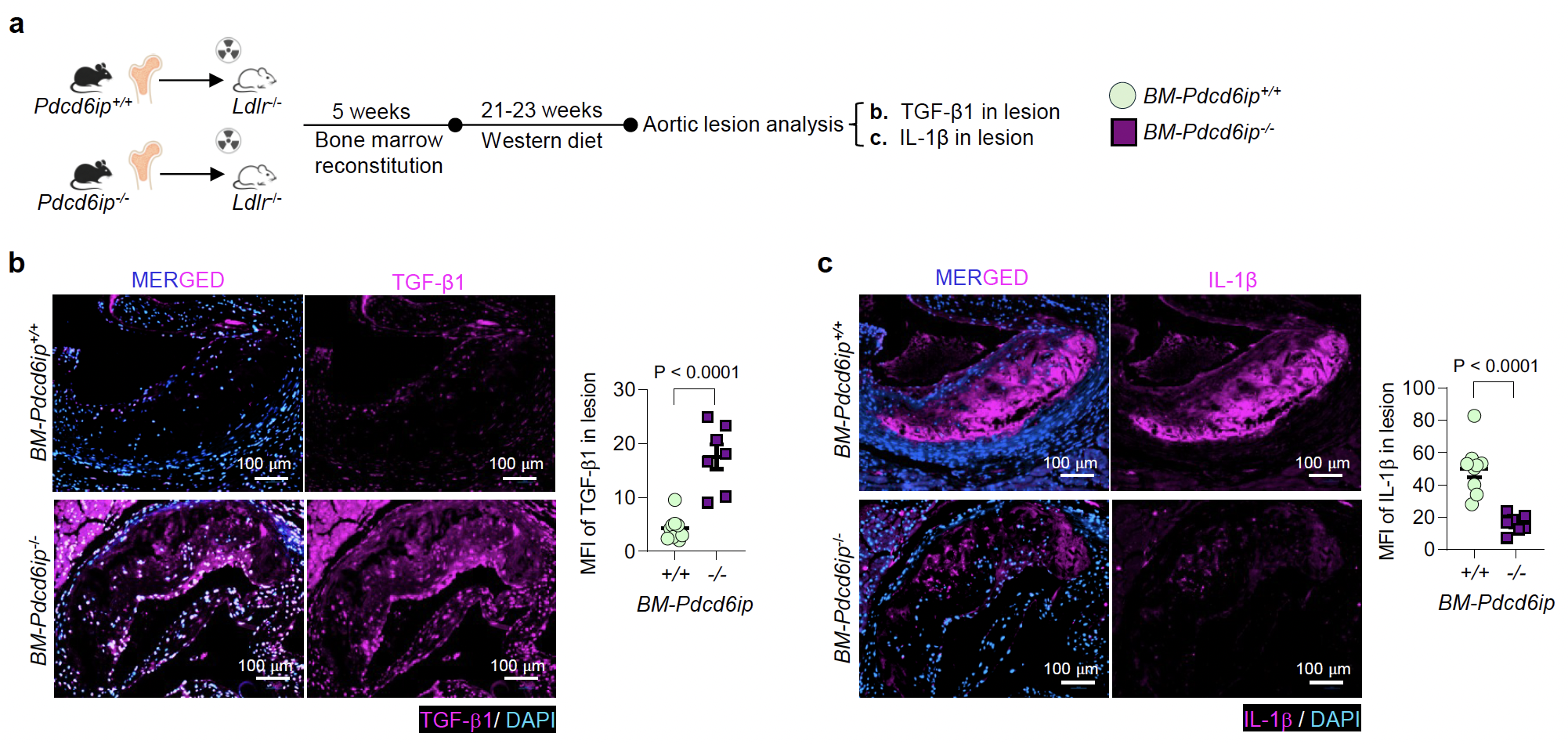

**Supplementary Fig. 4 | Hematopoietic *Pdcd6ip* deletion is associated with a less inflammatory, more resolution-associated plaque environment.**

**a**, Schematic of the study design, as in **Fig. 4**. *Ldlr*^−/−^ recipient mice were lethally irradiated and transplanted with bone marrow from *Pdcd6ip^+/+^* or *Pdcd6ip^−/−^* donor mice. After 5 weeks of reconstitution, mice were fed a Western diet for 21-23 weeks. Throughout this figure, *BM-Pdcd6ip^+/+^* and *BM-Pdcd6ip^−/−^* indicate *Ldlr^−/−^* recipient mice reconstituted with *Pdcd6ip^+/+^* or *Pdcd6ip^−/−^* bone marrow, respectively. **b,** Quantification of lesional TGF-β1 immunofluorescence staining, showing increased expression of the anti-inflammatory, resolution-associated cytokine TGF-β1 in plaques of *BM-Pdcd6ip^−/−^* compared with *BM-Pdcd6ip^+/+^* mice. *n* = 10 *BM-Pdcd6ip^+/+^* and 7 *BM-Pdcd6ip^−/−^* mice. **b**, Quantification of lesional IL-1β immunofluorescence staining, showing reduced expression of the pro-inflammatory cytokine IL-1β in plaques of *BM-Pdcd6ip^−/−^* compared with *BM-Pdcd6ip^+/+^* mice. *n* = 9 *BM-Pdcd6ip^+/+^* and 7 *BM-Pdcd6ip^−/−^* mice. Data are shown as mean ± SEM. Statistical significance was determined using a two-tailed Student’s *t*-test.

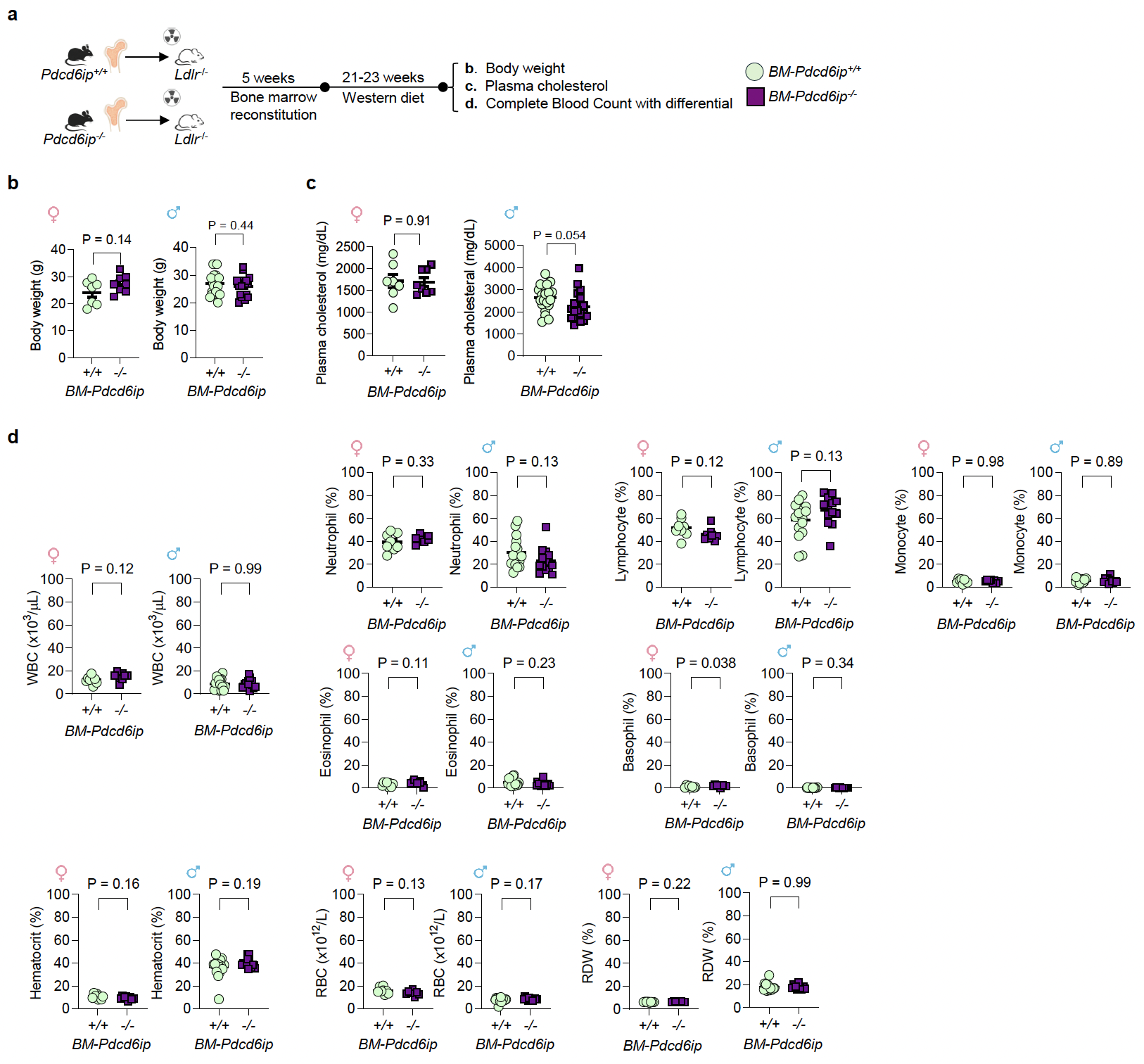

**Supplementary Fig. 5 | Hematopoietic *Pdcd6ip* deletion does not substantially alter body weight, systemic lipid levels, or circulating blood cell abundance in atherosclerotic mice.**

**a,** Schematic of the experimental design, as in **Fig. 4**. Body weight, plasma cholesterol level, and Complete Blood Count (CBC) with differential were assessed after 21-23 weeks of Western diet feeding. **b**, Body weight. Female: *n* = 7 *BM-Pdcd6ip^+/+^* and 8 *BM-Pdcd6ip^−/−^* mice; male: *n* = 20 *BM-Pdcd6ip^+/+^* and 18 *BM-Pdcd6ip^−/−^* mice. **c**, Plasma cholesterol levels. Female: *n* = 7 *BM-Pdcd6ip^+/+^* and 8 *BM-Pdcd6ip^−/−^* mice; male: *n* = 20 *BM-Pdcd6ip^+/+^* and 18 *BM-Pdcd6ip^−/−^* mice. **d**, CBC with differential. WBC, white blood cell; RBC, red blood cell; RDW, red blood cell distribution width. Female: *n* = 8 *BM-Pdcd6ip^+/+^* and 8 *BM-Pdcd6ip^−/−^* mice; male: *n* = 16 *BM-Pdcd6ip^+/+^* and 13 *BM-Pdcd6ip^−/−^* mice. Data are shown as mean ± SEM. Statistical significance was determined using a two-tailed Student’s *t*-test.

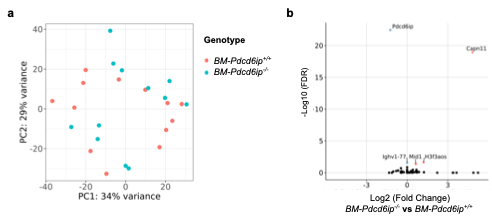
 **Supplementary Fig. 6 | Whole-spleen transcriptomic profiling of *Pdcd6ip^+/+^* and *Pdcd6ip^−/−^* bone marrow recipient mice.**

Bulk RNA-seq was performed on whole spleens from *BM-Pdcd6ip^+/+^* (n = 13) and *BM-Pdcd6ip^−/−^* (n = 12) recipient mice using paired-end sequencing, with 21–42 million reads per sample (mean = 31.6 million reads). **a,** Principal component analysis (PCA) of whole-spleen transcriptomes. **b,** Volcano plot of differential gene expression between *BM-Pdcd6ip^−/−^* and *BM-Pdcd6ip^+/+^* recipient mice. Differentially expressed genes were defined by an absolute fold change ≥1.5 and FDR <0.05; four genes met these criteria. Complete differential expression analysis is provided in **Supplementary Table 6**.

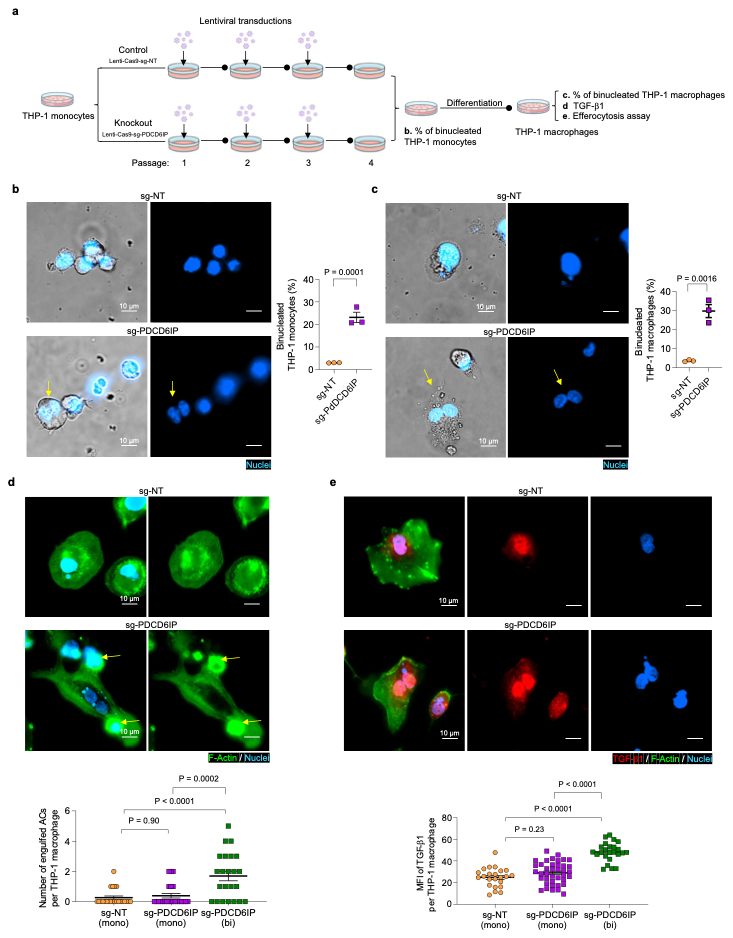

**Supplementary Fig. 7 | *PDCD6IP* perturbation generates binucleated, pro-efferocytic human macrophage-like cells.**

**a,** Schematic of the experimental design. Human THP-1 monocytes were transduced with lentiviral vectors encoding Cas9 together with *PDCD6IP*-targeting sgRNA (sg-PDCD6IP) or non-targeting control sgRNA (sg-NT). Cells were transduced immediately after passaging and allowed to expand before subsequent passaging and transduction, for a total of three rounds of transduction across three passages. Cells were analyzed after the third transduction/passaging cycle. Puromycin selection was initiated 48 h after the first transduction and maintained by refreshing puromycin-containing medium. This strategy enriched *PDCD6IP*-perturbed THP-1 monocytes. **b**, Quantification of binucleated THP-1 monocytes by live-cell imaging after Hoechst nuclear staining, showing that *PDCD6IP* perturbation increased the percentage of binucleated monocytes. Yellow arrows indicate binucleated THP-1 monocytes. *n* = 3 independent experiments. **c,** THP-1 monocytes after three rounds of lentiviral transduction were treated with 100 nM phorbol 12-myristate 13-acetate for 24h followed by 48 h culture without PMA to differentiate them into THP-1 macrophages, followed by Hoechst nuclear staining and live-cell imaging. Quantification showed that *PDCD6IP* perturbation increased the percentage of binucleated THP-1 macrophages. Yellow arrows indicate binucleated THP-1 macrophages. *n* = 3 independent experiments. **d**, Quantification of *in vitro* efferocytosis in THP-1 macrophages, showing enhanced efferocytosis capacity in *PDCD6IP*-perturbed binucleated (bi) macrophages, but not sg-PDCD6IP targeted mononuclear (mono) macrophages. THP-1 macrophages were incubated with ACs at a 1:1 ratio for 45 min, fixed with 4% formaldehyde, and stained for F-actin (Phalloidin) and nuclei (Hoechst). Yellow arrows indicate engulfed ACs by THP-1 macrophages. *n* = 23 cells per group from 3 independent experiments. **e**, Quantification of TGF-β1 expression in THP-1 macrophages, showing increased TGF-β1 levels in *PDCD6IP*-perturbed binucleated macrophages. THP-1 macrophages were immunostained with anti-TGF-β1 antibodies and co-stained with F-actin (Phalloidin) and nuclei (Hoechst). *n* = 23 cells from sg-NT (mono), 39 cells from sg-PDCD6IP (mono), and 24 cells from sg-PDCD6IP (bi) from 3 independent experiments. Data are shown as mean ± SEM. All images were acquired by Nikon Ti-S Automated Inverted Microscope equipped with a Plan Apo λ 60×/1.40 oil objective. Statistical significance was determined using a two-tailed Student’s t-test for **b** and **c**, and one-way ANOVA with Tukey’s post hoc test for **d** and **e**.
